# Pharmacological Enhancement of Adult Hippocampal Neurogenesis Improves Behavioral Pattern Separation in Young and Aged Mice

**DOI:** 10.1101/2024.02.01.578406

**Authors:** Wei-li Chang, Karly Tegang, Benjamin A. Samuels, Michael Saxe, Juergen Wichmann, Denis J. David, Indira Mendez David, Angélique Augustin, Holger Fischer, Sabrina Golling, Jens Lamerz, Doris Roth, Martin Graf, Sannah Zoffmann, Luca Santarelli, Ravi Jagasia, René Hen

## Abstract

**BACKGROUND:** Impairments in behavioral pattern separation (BPS)—the ability to distinguish between similar contexts or experiences—contribute to memory interference and overgeneralization seen in many neuropsychiatric conditions, including depression, anxiety, PTSD, dementia, and age-related cognitive decline. While BPS relies on the dentate gyrus and is sensitive to changes in adult hippocampal neurogenesis (AHN), its significance as a pharmacological target has not been tested.

**METHODS:** In this study, we applied a human neural stem cell high-throughput screening cascade to identify compounds that increase human neurogenesis. One compound with a favorable profile, RO6871135, was then tested in BPS in mice.

**RESULTS:** Chronic treatment with RO6871135, 7.5 mg/kg increased AHN and improved BPS in a fear discrimination task in both young and aged mice. RO6871135 treatment also lowered innate anxiety-like behavior, which was more apparent in mice exposed to chronic corticosterone. Ablation of AHN by hippocampal irradiation supported a neurogenesis-dependent mechanism for RO6871135-induced improvements in BPS. To identify possible mechanisms of action, in vitro and in vivo kinase inhibition and chemical proteomics assays were performed. These tests indicated that RO6871135 inhibited CDK8, CDK11, CaMK2a, CaMK2b, MAP2K6, and GSK3b. An analog compound also demonstrated high affinity for CDK8, CaMK2a, and GSK3b.

**CONCLUSIONS:** These studies demonstrate a method for empirical identification and preclinical testing of novel neurogenic compounds that can improve BPS, and points to possible novel mechanisms that can be interrogated for the development of new therapies to improve specific endophenotypes such as impaired BPS.

## Introduction

Pattern separation is the process of separating overlapping sensory information, contexts, and experiences, into distinct neural representations. It is believed that this process facilitates the rapid storage of new memories without inducing large amounts of interference (1–3). Computational theories and simulations predicted that this role is performed by the dentate gyrus (DG) (4–7). This function of the DG was later empirically established in rodents (8–14). In mammals, the DG is one of two brain regions (with the subventricular zone), that continues to generate new neurons throughout development and adulthood, a phenomenon known as adult hippocampal neurogenesis (AHN)(15,16).

Previous work from our group and others has shown the importance of AHN for behavioral measures of pattern separation (BPS): ablating AHN in mice causes impairments in BPS, while enhancing AHN with exercise or enrichment improves it (17–25). A genetic manipulation specifically targeting AHN was sufficient to improve BPS in mice, as measured by a fear discrimination task, underscoring the role of new neurons in this cognitive task (25). In addition, transiently silencing immature adult-born granule cells during discrete epochs of a fear discrimination task can disrupt pattern separation (26). Stress has been shown to decrease AHN (27–34), and there is evidence that enhanced AHN confers increased resilience to chronic stress (35–39). AHN also decreases dramatically with age (18,40–43), as does BPS performance (20,44–46).

In humans, tasks have been developed to test pattern separation, and when studied in conjunction with fMRI, they have also been shown to reliably engage the DG and downstream CA3 region (14,47–55). Deficits in BPS may contribute to overgeneralization of negative emotion seen in depression, anxiety, and trauma-related disorders (56–62). Pattern separation also declines with aging in humans (55,63,64), an effect that is even more pronounced in patients with mild cognitive impairment (50,55,65) and further impaired in Alzheimer’s disease (65,66).

Based on these observations, enhancement of AHN is thought to be a promising target for therapeutic development to treat conditions demonstrating BPS deficits, such as depression, anxiety disorders, PTSD, and age-related cognitive decline as well as dementia. In the present study, a high-throughput in vitro screening cascade was used to empirically identify compounds with human neurogenic properties. One family of promising neurogenic molecules from this screen, piperazinones, was chemically optimized and the resulting compound RO6871135 was then tested in vivo. We found that RO6871135 enhanced AHN and improved BPS in a neurogenesis-dependent manner.

## Methods and Materials

See the Supplement for detailed methods and materials.

### High-throughput screen for human neurogenesis

Human neural stem/progenitor cells (hNSCs) were derived from human embryonic stem cells (hESCs) according to previously reported procedures (67,68).

### Animal Care

All experimental procedures were conducted in compliance with the NIH Guide for the Care and Use of Laboratory Animals and approved by the Institutional Animal Care and Use Committee at the New York State Psychiatric Institute. Chronic corticosterone experiments were conducted in compliance with protocols approved by another Institutional Animal Care and Use Committee (Council directive no. 87–848, 19 October 1987, Ministère de l’Agriculture et de la Forêt, Service Vétérinaire de la Santé et de la Protection Animale). Mice were housed two to five per cage and maintained on a 12 h light/dark schedule with access to food and water *ad libitum*, except when otherwise stated.

### Drug administration

For all studies with behavioral testing, adult male c57BL/6 mice received 7.5 mg/kg of RO6871135 daily by oral gavage for 21 days before behavioral testing. On the days of behavioral testing, animals were gavaged after behavior was completed.

### Behavior

Anxiety-related and BPS behavioral tasks were carried out as described in Supplemental Methods. For fear discrimination context parameters, see Tables S1 and S2. Hippocampal irradiation (69–71) and chronic corticosterone (72) interventions were performed as described previously.

### Binding analyses

In vitro pharmacological screening for off-target effects was also performed as described (73), tested at Cerep (now Eurofins Pharma Discovery). In vitro kinase screening assays were performed to determine kinase activity inhibition as described (74) via LeadHunter® Drug Discovery Services Panels (Eurofins DiscoverX Products, LLC, Fremont, CA), and dissociation constants (K_d_) for compound-kinase interactions were calculated. In situ kinase binding was assayed using the KiNativ^TM^ platform (75–77), (ActivX, La Jolla, CA). Brain and liver tissue samples were collected from RO6871135- or vehicle-treated mice. Chemical proteomics analysis was conducted in hNSCs using two close chemical analogues of RO6871135: one neurogenically activity and one neurogenically inactive compound.

### Statistical analyses

Statistical analyses were carried out using the Python packages statsmodels (78) and SciPy (79), R 4.3.2, and GraphPad Prism (version 9.5.1 for macOS, GraphPad Software, San Diego, California, USA, www.graphpad.com.) For all comparisons, values of *p <* .05 were considered as significant.

## Results

### In vitro neurogenesis screen

Neural stem cells (hNSCs) were derived from human embryonic stem cells (hESCs) as previously described (67,80) (Figure S1A). hNSCs were exposed to factors known to modulate neurogenesis, then stained for DAPI to quantify cell number, and Tuj1, a marker of immature neurons (Figure S1B). As expected, DAPT, which blocks Notch signaling, accelerated differentiation (81), resulting in upregulation of Tuj1 and reduced proliferation, evidenced by reduced cell number. Consistent with previous in vitro and in vivo findings, Wnt-3a promoted both proliferation and differentiation of hNSCs to immature neurons (82,83). The addition of the known mitogen fibroblast growth factor FGF-2 also promoted proliferation of neural progenitor cells (NPCs) while inhibiting neuronal differentiation. Recapitulation of these known effects supported the use of hNSCs as a model to screen for novel human neurogenic modulators (Figure S2A). ∼1 million compounds were screened, and the results are plotted as a histogram with >10,000 and >3,000 hits, 3 and 4 standard deviations from the mean, respectively (Figure S2B). The dose-response curve of a chemically optimized molecule from an original hit, RO6871135 (Figure 1A), had a potency of 26 nM for increasing hNSC cell counts (measured by quantifying ATP as an indicator of metabolically active cells), and was essentially inactive on the counter screen in hESC-derived mesenchymal stem cells (hMSCs, Figure S2C). To determine if RO6871135 was truly neurogenic, high content screening was performed quantifying both nuclei (DAPI stain) and immature neurons (Tuj1 stain, Figure 1B). Dose-response curves of RO6871135 on cell number (Figure S2D) and neurite network (Figure S2E) revealed potency in the range of 20 nM, in line with the potency seen in the previous step of the screen, which used an ATP assay to quantify cell number. RO6871135 was profiled in a standard battery of drug development assays at Roche (73), such as hepatic enzyme activity effects and other safety tests, and this compound had a favorable profile (Table S3). Off-target assays to assess risk for adverse drug reactions (73,84) indicated no pharmacological activity at concentrations relevant to in vitro potencies and in vivo testing (Table S4).

**Figure 1:**
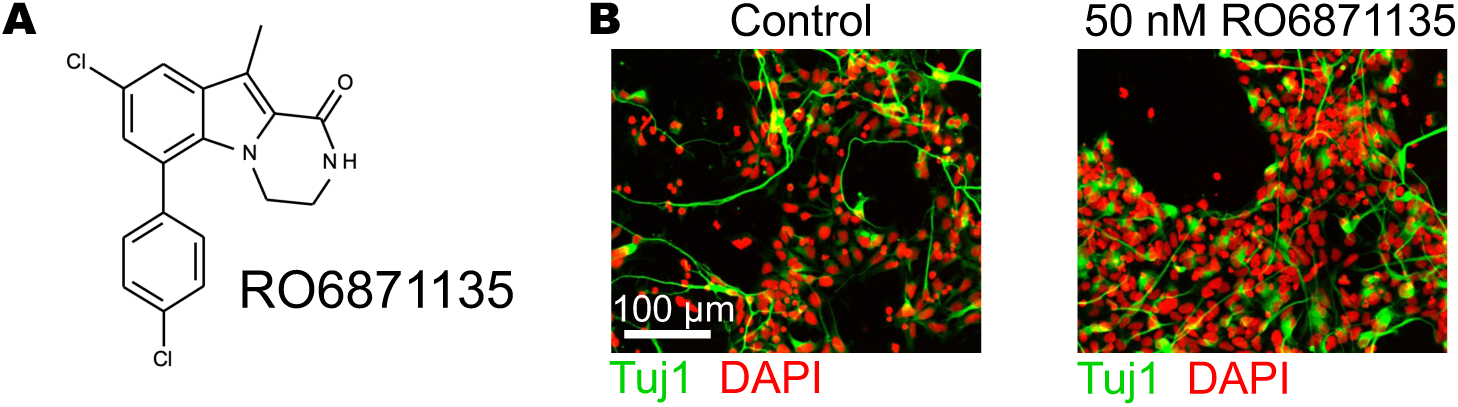
RO6871135 increases in vitro human neurogenesis. (A) Molecular structure of RO6871135. (B) Representative images of differentiating human embryonic stem cell derived neural stem cells (hNSCs) in the presence or absence of 50nM of RO6871135 in the media. DAPI in red for cell number, including hNSCs and neural progenitor cells (NPCs), while Tuj1 staining is shown in green reflects initial differentiation or immature neurons.

### In vivo screening for increased neurogenesis

RO6871135 showed good pharmacokinetic parameters after single-dose administration in mice (Table S3). After 14 days of PO administration of the compound (Figure 2A top), there were dose-dependent increases in markers of proliferation (as measured by Ki67 staining, Figure 2B left). Using a BrdU paradigm to assess survival of adult-born cells, there was a dose-dependent increase in BrdU staining (Figure 2B middle), and increased numbers of immature neurons (as measured by DCX staining, Figure 2B right). For behavioral experiments, a longer treatment schedule was used prior to testing to allow for the accumulation of immature granule cells, which generally require at least 2 weeks to begin integrating into the surrounding circuit (85,86).

**Figure 2:**
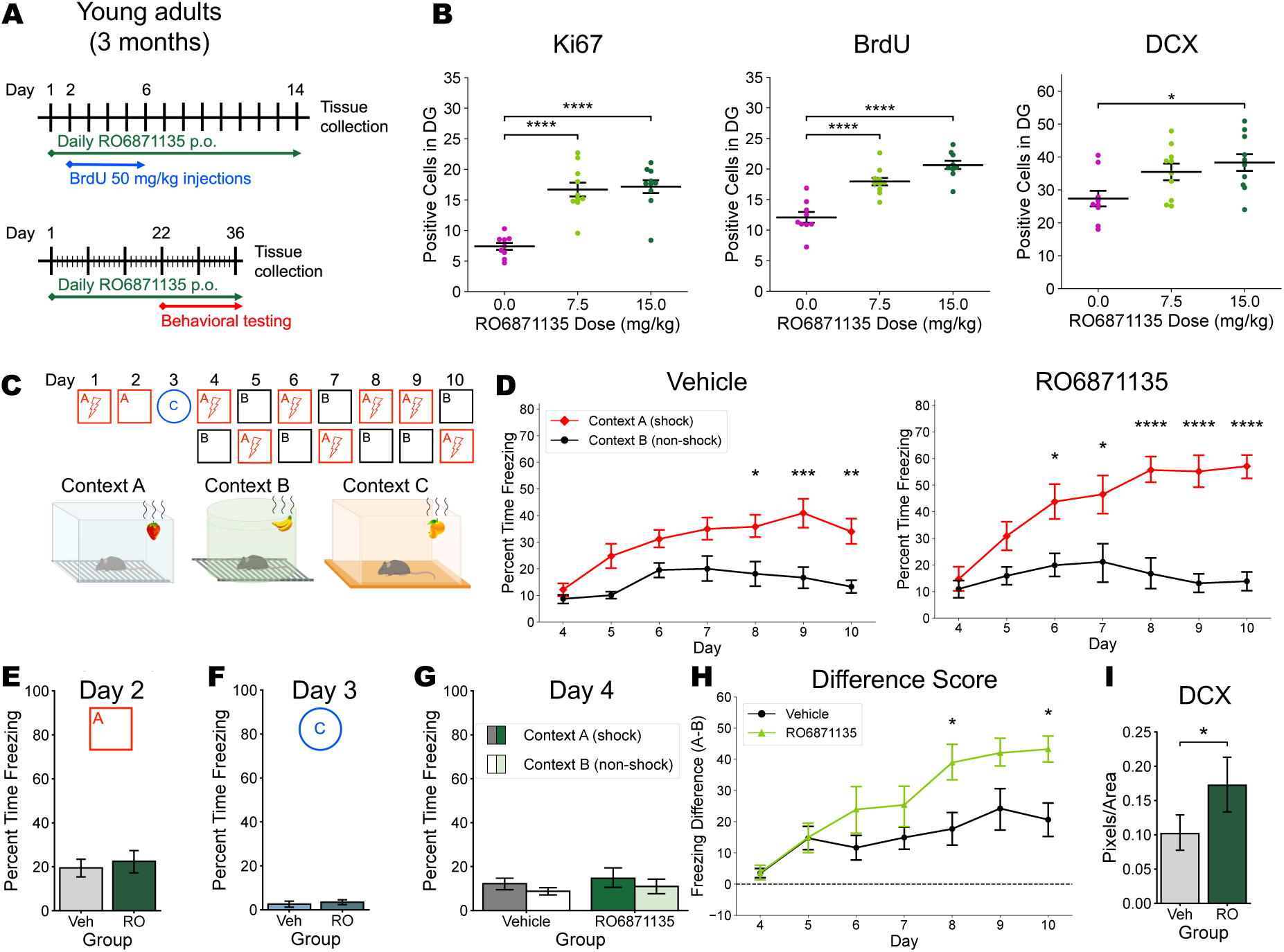
Chronic in vivo administration of RO6871135 increases neurogenesis and improves pattern separation. (A) Timelines of experimental design. 14-day administration for histology studies is shown on the top row. 21-day administration for behavioral testing followed by histology shown on the bottom row. (B) Positive cell counts (±SEM) for Ki67, BrdU, and DCX. There was a significant effect of treatment group in all measures (Ki67 F_(2,26)_= 26.64, *p*<0.0001; BrdU F_(2,25)_=30.60, *p*<0.0001; DCX F_(2,26)_= 4.658, *p*<0.05). Controls (*n* = 9) had lower cell counts of Ki67 (*p*<0.0001) and BrdU (*p*<0.0001) compared to mice treated with 7.5 mg/kg of RO6871135 (*n* = 10), and they had lower cell counts of all three markers compared to mice treated with 15 mg/kg of RO6871135 (n=10; Ki67 *p*<0.0001; BrdU *p*<0.0001; DCX *p*<0.05). (C) Schematic for contextual fear conditioning and fear discrimination tasks. (D) Percent time freezing across days in the fear discrimination task. Vehicle-treated controls (*n* = 8) had a significant context × day interaction (F_(6,84)_= 2.876, *p*<0.05), with significant differences in freezing starting on Day 8. RO6871135-treated mice (*n* = 8) also had a significant context × day interaction (F_(6,84)_= 7.899, *p* <0.0001), with significant differences in freezing between contexts by Day 6. (E-F) Freezing time after single-shock contextual fear conditioning in the same context or a novel context. RO6871135 treatment did not alter expression of contextual fear (*t*(14)= −0.434, NS) and did not affect generalization of fear to a different, novel context (*t*(14)= −0.490, NS). (G) Freezing time on the first day of exposure to the similar Context B at the beginning of the fear discrimination task, showing similar freezing levels to the shock Context A in both groups (context F_(1,28)_= 1.157, NS; group F_(1,28)_= 0.500, NS; group × context F_(1,28)_= 0.001, NS). (H) Difference score across days, calculated by subtracting the freezing time in Context B from freezing time in Context A. There was a significant effect of group (F_(1,14)_= 5.948, *p<*0.05) and a significant group × day interaction (F_(6,84)_= 2.256, *p<*0.05). Post-hoc testing indicated significant differences on Days 8 and 10 (*p<*0.05). (I) Doublecortin staining in dentate sections from the same mice that underwent behavioral testing. After over 5 weeks of treatment with 7.5 mg/kg of RO6871135, there is a significant increase in DCX staining (*t*(15) = −2.648, *p<*0.05). *p*≥0.05 is not significant, \**p*<0.05, \*\**p*<0.01, \*\*\**p*<0.001, \*\*\*\**p*<0.0001. p.o., per oral; DCX, doublecortin; RO, RO6871135.

### Chronic administration of RO6871135 alters contextual fear discrimination for similar contexts without effecting contextual fear conditioning

After >21 days of treatment with RO6871135 (Figure 2A bottom), there was no observed difference in the level of freezing on the retrieval day of contextual fear conditioning (Figure 2E). There was also negligible freezing in both groups in a novel context on Day 3 (Figure 2F). On Day 4, when mice were re-exposed to Context A and given a foot shock, followed by exploration of the similar Context B, both groups exhibited comparable levels of freezing between the two contexts (Figure 2D, G). On subsequent days of the fear discrimination task, freezing levels in Context A increased while freezing in Context B remained relatively stable (Figure 2D). Repeated measures (rm) ANOVA of freezing in vehicle-treated mice indicated a significant effect of context and day, and a significant context × day interaction. Mice in the vehicle-treated group successfully discriminated between the two contexts, as measured by freezing time, starting on Day 8, or the fifth day of exposure to both contexts. In mice treated with RO6871135, this discrimination was achieved by Day 6, or the third exposure to both contexts. The drug-treated mice were thus able to discriminate between similar contexts two days faster than vehicle-treated mice. Freezing differences between Contexts A and B were calculated for each mouse, and these difference scores were compared between experimental groups with rm ANOVA (Figure 2H). There was a significant effect of group, day, and a group × day interaction; Days 8 and 10 were significantly different by post hoc testing. We confirmed that there was increased doublecortin (DCX) staining after RO6871135 in these same mice (Figure 2I).

### Aged mice have lower levels of AHN and a deficit in BPS compared to young mice; these effects are partially rescued with RO6871135

We tested RO6871135 in aged mice (>18 months) using the same drug administration schedule (Figure 3A), followed by behavioral testing and immunohistochemistry. Compared to young mice in the same experimental cohort, we saw a dramatic decrease in all measures of neurogenesis in vehicle-treated aged mice, in keeping with previous findings (18,40–43). RO6871135 in aged mice significantly increased detectable BrdU (administered 4 weeks before tissue collection), and DCX staining (Figure 3B). For the fear discrimination task, we used a non-randomized order for context presentation (Figure 3C), since aged mice were expected to have difficulty with the randomized order paradigm (20). Young and aged mice showed comparable levels of freezing in the shock-associated Context A after a single day of exposure, indicating no effect of age on contextual fear conditioning (Figure 3D). There was no significant difference in freezing between Contexts A and B in young mice on the first day of exposure to both contexts, but significantly higher freezing levels in Context B in both vehicle- and RO6871135-treated aged mice (Figure 3D). By the next day, young mice were already discriminating between the two contexts, while vehicle- and RO6871135-treated aged mice had nearly identical freezing levels in Contexts A and B (Figure 3E). Vehicle-treated aged mice failed to discriminate between contexts during the initial comparison period (through Day 9); after two more days of exposure, they did eventually discriminate (Figure S3). RO6871135 treatment of aged mice improved BPS, with successful discrimination after Day 5 (Figure 3E). We did not observe an effect of RO6871135 treatment in young-adult mice using this behavioral paradigm, since vehicle-treated young mice already discriminated by the second day of exposure to both contexts (Figure S4).

**Figure 3:**
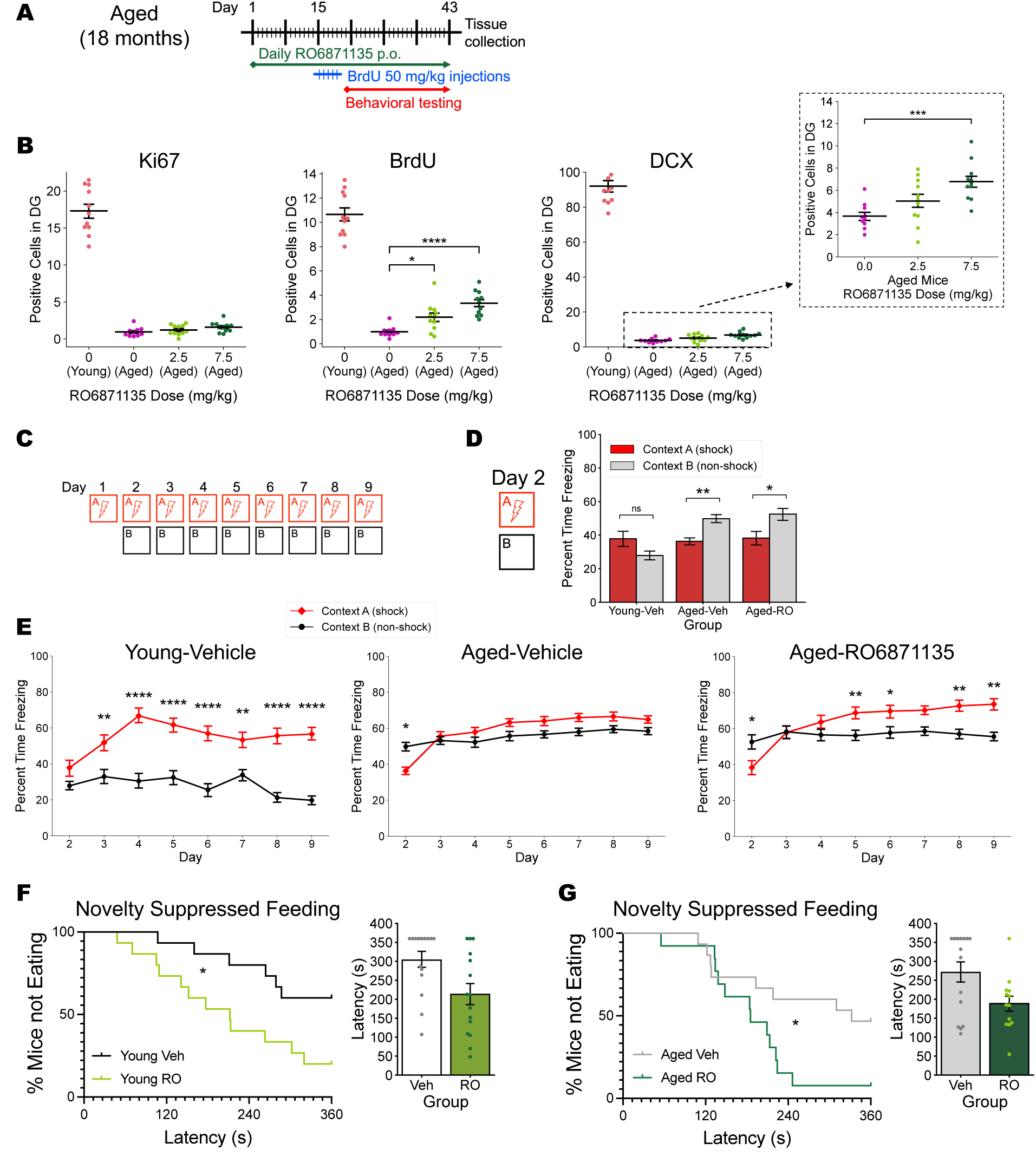
Aged mice have much lower measures of AHN. They also have a reduced fear discrimination, which is partially rescued with treatment by RO6871135. (A) Timeline of experimental design. The same cohort underwent behavioral testing and histological analysis. (B) Positive cell counts (±SEM) for Ki67, BrdU, and DCX. Measures from vehicle-treated young mice (*n* = 11) are shown for reference, compared to vehicle-treated aged mice (*n* = 10). Among aged mice, RO6871135 significantly increases counts of cells positive for BrdU (F_(2,29)_=15.99, *p*<0.0001) and DCX (F_(2,29)_=8.745, *p*<0.005) but does not restore them to the level of young mice. (C) Schematic for non-randomized fear discrimination task. (D) Freezing time after single-shock contextual fear conditioning in the same context A one day after foot shock or on first exposure to the similar context B (context F_(1,128)_= 3.672, NS; group F_(2,128)_= 6.091, *p*<0.005; group × context F_(2,128)_= 6.754, *p*<0.005). There was no significant effect of context in the young mice and elevated freezing in the similar context B in both groups of aged mice (vehicle *p*<0.005; RO6871135 *p*<0.05). (E) Percent time freezing across days in the fear discrimination task in young and aged mice. Vehicle-treated young mice had a significant context × day interaction (F_(7,196)_= 4.598, *p<*0.0001), with significantly higher freezing in the shock context by the second day of exposure to both contexts. Vehicle-treated aged mice also had a significant context × day interaction (F_(7,378)_= 8.994, *p<*0.0001), but beyond elevated freezing in Context B on the first day, did not demonstrate differential freezing across the subsequent 7 days. Aged mice after treatment with RO6871135 had a significant context × day interaction (F_(7,322)_= 9.860, *p<*0.0001), again with elevated freezing in Context B on Day 2, but with significantly higher freezing in the shock context on Days 5, 6, 8, and 9 of the experiment. (F) Latency to feed in the NSF tests in young and aged (G) mice, represented as a survival curve on the left and as the latency measures on the right. Log-rank (Mantel-Cox) test: Young Chi square= 5.680, *p*<0.05; Aged Chi square=4.854, *p*<0.05. There was no significant effect of RO6871135 on latency to feed in the home cage in either age group (data not shown). *p*≥0.05 is not significant, \**p*<0.05, \*\**p*<0.01, \*\*\**p*<0.001, \*\*\*\**p*<0.0001. p.o., per oral; DCX, doublecortin; RO, RO6871135.

### RO6871135 alters innate anxiety-like behavior in the novelty suppressed feeding (NSF) test. Aging alters behavior in the OFT

In the NSF test, which measures approach-avoidance behavior in the latency to feed in a novel arena, treatment with RO6871135 significantly decreased latency to feed in both young (Figure 3F) and aged mice (Figure 3G). However, in another innate anxiety-like measure, the open field test (OFT), mice treated with RO6871135 showed no difference compared to vehicle-treated mice in exploration of the center zone (Figure S5B, S5C); there was increased locomotor activity (Figure S5A).

Aged mice exhibited lower levels of locomotion in the OFT, and RO6871135 increased locomotion in both age groups with no significant age × treatment interaction (Figure S5A). Aged mice spent more time and travelled more in the center zone in the setting of overall lower locomotor activity, and we did not observe a differential effect of RO6871135 on OFT measures in young vs. aged mice (Figure S5B, S5C). When comparing latency to feed in the NSF test between vehicle-treated young and aged mice, there was no difference between the age groups (Figure S6).

### Permanent ablation of neurogenesis by irradiation blocks the effects of RO6871135 on contextual fear discrimination but not latency to feed in NSF

To investigate whether immature granule cells were required for the effects of RO6871135 on pattern separation, we used a well-established method of bilateral X-irradiation to permanently ablate AHN (69–71), followed by two months of recovery from inflammatory effects of irradiation before initiation of treatment with RO6871135 (Figure 4A). Animals were then tested with the same randomized-order fear discrimination task used in young, non-irradiated mice to assess drug effects in the absence of AHN. Levels of freezing after contextual fear conditioning and on the first exposure to both similar contexts were comparable in irradiated vs. non-irradiated mice, and there was no effect of drug treatment on these days (Figure S7). We saw a decrease in BPS performance after irradiation relative to non-irradiated mice, with a significant context × day interaction in vehicle-treated irradiated mice, but no discrimination between contexts A and B until Day 10 (Figure 4B). Freezing in irradiated mice treated with RO6871135 showed no significant effect of context, and no context × day interaction (Figure 4C). The difference scores of freezing in A-B showed no significant effect of treatment group, and no group × day interaction (Figure 4D). Irradiation did not block the effect of RO6871135 in the NSF test, and drug-treated irradiated mice exhibited decreased latency to feed compared to vehicle-treated irradiated controls (Figure 4E). Ablation of AHN in irradiated mice was confirmed histologically after completion of behavioral testing (Figure 4F).

**Figure 4:**
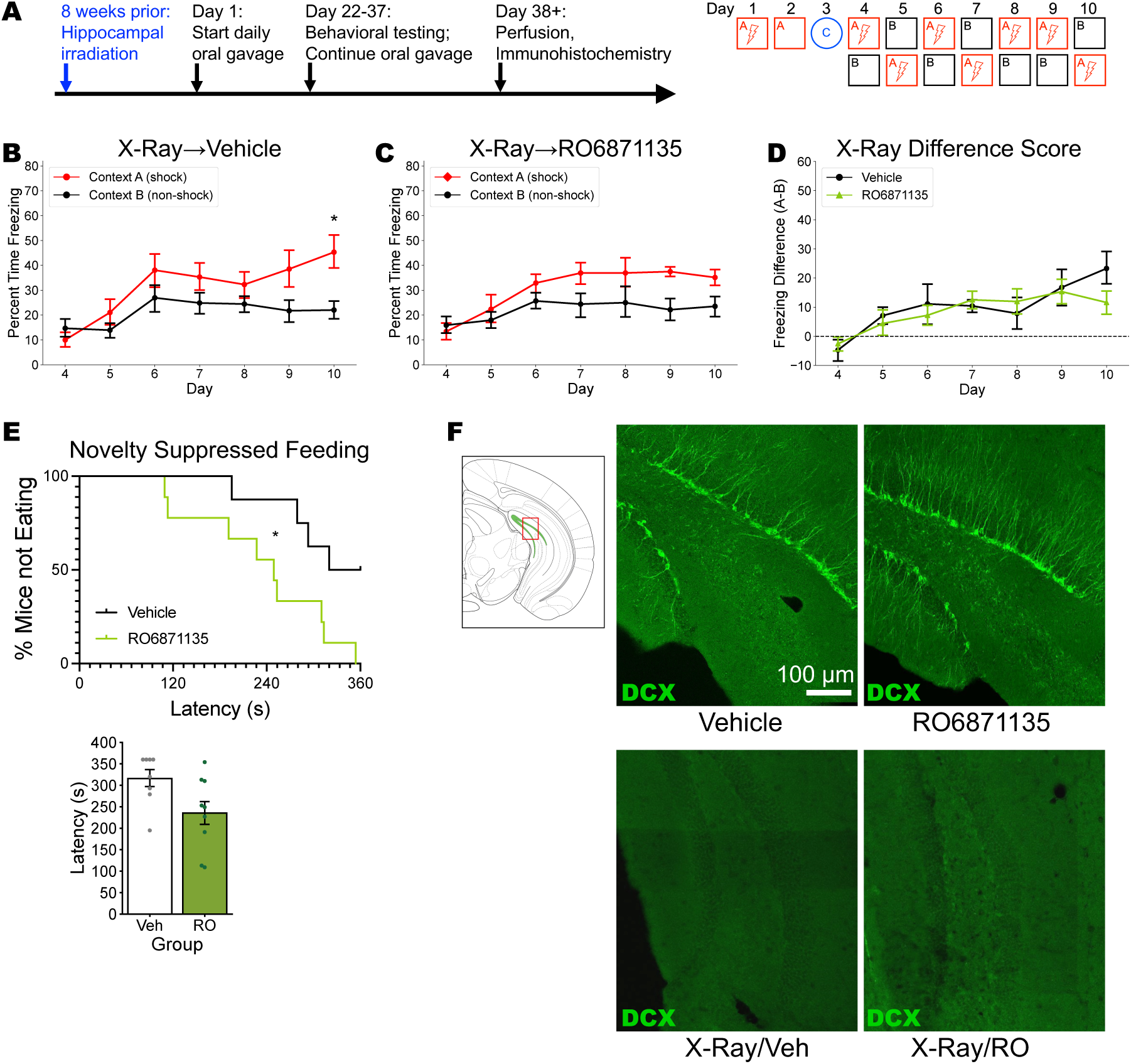
Irradiation blocks the effect of RO6871135 on behavioral pattern separation, but not on NSF latency to feed. (A) Timeline of focal irradiation followed by 8 weeks of recovery and then initiation of daily dosing of RO6871135 and behavioral testing, as conducted with non-irradiated mice. (B) Vehicle-treated mice with chronic ablation of adult hippocampal neurogenesis showed a significant context x day interaction, with a significant difference in freezing on Day 10 (*n* = 9). (C) Irradiated mice (*n* = 9) that received RO6871135 (*n* = 9) had no significant effect of context (F_(1,16)_= 3.333) and no context × day interaction (F_(6,96)_= 1.889). (D) The difference scores had a significant effect of day (F_(6,90)_= 6.607, *p* < 0.0001), but no significant effect of drug group (F_(1,15)_= 0.1623, NS) and no group × day interaction. (E) Latency to feed in the NSF test shows that decreased latency in the RO6871135-treated group remains even after irradiation. (Top) Survival curve log-rank (Mantel-Cox) test: Chi square = 6.034, *p*<0.05. (Bottom) Latency measures of individual mice in the NSF. There was no significant effect of RO6871135 on latency to feed in the home cage (data not shown). (F) Top left panel shows an atlas image of the dentate gyrus and approximate field of view for microscope images (red rectangle). Following panels show representative images of doublecortin staining in non-irradiated and irradiated mice treated with vehicle of RO6871135. Lack of staining in irradiated mice confirms ablation of AHN and lack of immature neurons observable by DCX staining. \**p*<0.05. DCX, doublecortin.

### RO6871135 reverses behavioral effects of chronic corticosterone and stimulates AHN

We next looked at RO6871135 effects after chronic corticosterone exposure (Figure 5A), a model of chronic stress used for anxiety- and depression-related models. Chronic corticosterone increased innate anxiety-like behavior, as measured by exploration of the center zone, and these effects were reversed in mice that had received RO6871135 (Figure 5B). Corticosterone exposure also increased anxiety-like behavior in the NSF test, with increased latency to feed. Treatment with RO6871135 partially reversed this effect and significantly decreased latency to feed compared to the corticosterone/vehicle group (Figure 5C). RO6871135 increased the number of DCX-positive cells in the setting of chronic corticosterone treatment as well (Figure 5D, E).

**Figure 5:**
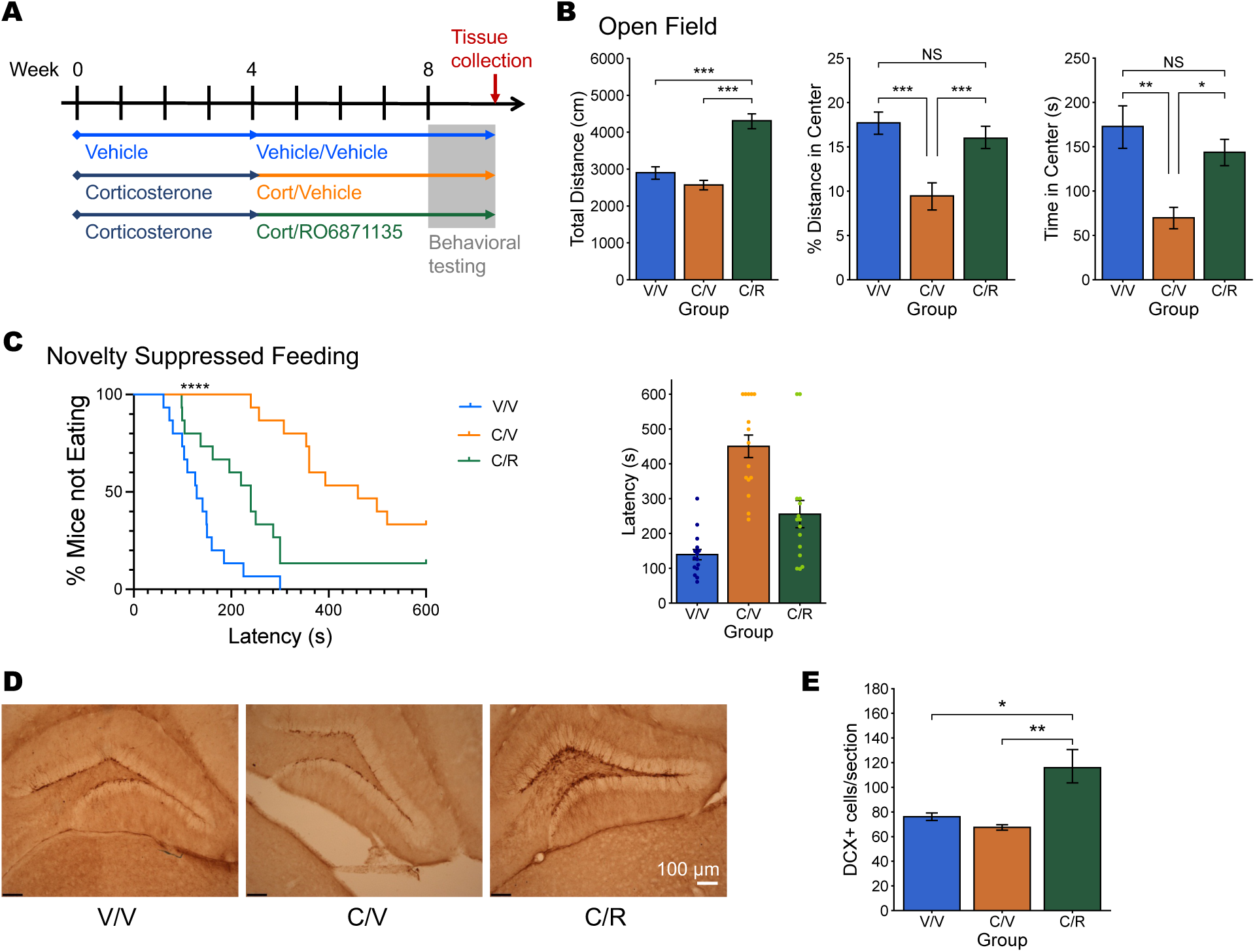
Chronic corticosterone increases anxiety-like behavior, which is reversed by treatment with RO6871135. (A) Experimental timeline. Mice are treated with vehicle or corticosterone for 4 weeks before starting daily RO6871135 or vehicle treatment for another 4 weeks while corticosterone or vehicle is continued. (B) There was no significant effect of corticosterone on total distance traveled in an open field, but RO6871135 increased locomotion compared to both the vehicle and corticosterone groups (F_(2,42)_=26.828, *p*<0.0001) Stars indicate significant differences in post hoc testing by Tukey’s HSD Test for multiple comparisons. Chronic corticosterone decreased the percent of distance traveled in the center of the open field arena, and this effect was reversed with RO6871135 treatment (F_(2,42)_=9.724, *p*<0.001). Mice that received chronic corticosterone also spent decreased time the center of the open field arena, and this effect was reversed with RO6871135 treatment (F_(2,42)_=8.421, *p*<0.001). (C) Chronic corticosterone increased latency to feed in the NSF test, and RO6871135 partially reversed this effect, represented as a survival curve and the latency values per mouse. Log-rank (Mantel-Cox) test: Chi square = 32.98, *p*<0.0001. Bonferroni-corrected α=0.017 for multiple comparisons, and all three treatment groups had significantly different latency values compared to either other group. There was no difference in home cage food consumption relative to body weight between vehicle/vehicle mice and those treated with RO6871135 (data not shown). (D) Representative images of doublecortin (DCX) staining in mice treated with vehicle vs. corticosterone and vehicle vs. RO6871135. (E) Quantification of DCX staining (F_(2,18)_=8.297, *p*<0.01). There was no significant change in DCX staining from chronic corticosterone alone, but RO6871135 treatment increased DCX in corticosterone-exposed mice. *p*≥0.05 is not significant, \**p*<0.05, \*\**p*<0.01, \*\*\**p*<0.001, \*\*\*\**p*<0.0001. V/V= vehicle/vehicle, C/V= corticosterone/vehicle. C/R= corticosterone/RO6871135.

### Functional activity and binding profiles of RO6871135

As stated above, a panel of assays to screen for GPCR binding was negative at relevant concentrations (Table S4). In order to identify putative targets of RO6871135 that could inform its mechanism of action, a series of binding assays were performed. In vitro binding against a panel of 96 kinases with the KINOMEscan^TM^ panel (Table S5) revealed significant functional inhibitory activity for CDK11. Though not included in the initial inhibition assay, CDK8 was added for calculation of K_d_ values. Inhibitory activity for CDK8 and CDK11, but no other kinases, was seen at sub-micromolar concentrations (Table S6).

Following the biochemical assays above, we then tested for activity in murine brain tissue. In situ kinase profiling was performed using KiNativ^TM^ for brain tissue from RO6871135-treated mice (Tables S7, S8). Liver tissue from the same animals was used for comparison (Table S8). Based on prior validation (75), >35% inhibition was considered significant. RO6871135 caused >50% inhibition for CDK8 and CDK11, as well as for CaMK2a, CaMK2b, and MAP2K6 in the brain.

In addition, a chemical proteomics study was conducted to define the potential targets of RO6871135 in hNSCs (Figure S8). The enriched proteins on the active RO6871135 analog versus the inactive one are highlighted in Figure S9. CaMK2a is, among the statistically significant differences, the most enriched protein on the active vs. inactive analogs. Seven kinases (GSK3A, GSK3B; MAPK1; MAPK3; CAMK2G, CSNK1A1 and CDK8) exhibited more binding to the active RO6871135 analog than the inactive one.

Top hits from in vitro (KINOMEscan^TM^), in situ (KiNativ^TM^), and chemical proteomics assays are summarized in Table 1. CDK8 activity was found in all three assays. CDK11 and GSK3B were among to top candidates in two out of three assays. CaMK2A, which was not directly tested in the KINOMEscan^TM^ assay, was strongly positive in the other two assays. CaMK2B, and MAP2K6, were also not directly tested in the KINOMEscan^TM^ assay, but were strongly positive in the in situ assay.

**Table 1:** A summary of proteins that showed significant binding or activity changes in the presence of RO6871135. KINOMEscan^TM^ results represent an in vitro assay of RO6871135. The KiNativ^TM^ assay was performed in situ on brain tissue collected from mice treated with RO6871135. CDK8, found across all three assays, is highlighted in yellow. Other proteins identified in two assays are highlighted in blue. Two proteins with high inhibition in the KiNativ^TM^ assay were not tested in the KINOMEscan^TM^ assay. + indicates a significant hit for that assay, while ++ denotes more binding or activity relative to the other positive hits. Raw values for individual assays are shown in Figure S9, and Tables S4-8.

| Assay: | KINOMEScan™ | KiNativ™ | Chemoproteomics |
| --- | --- | --- | --- |
| CDK11 | ++ | ++ |  |
| CDK8 | + | ++ | + |
| CaMK2A | Not tested | ++ | ++ |
| CaMK2B | Not tested | ++ |  |
| GSK3A |  |  | ++ |
| GSK3B |  | + | ++ |
| MAP2K6 | Not tested | ++ |  |

## Discussion

We demonstrate that treatment with RO6871135 was sufficient to enhance AHN and improve BPS in both young and aged mice, and that BPS effects are neurogenesis-dependent. This candidate compound was identified from a screening cascade testing for neurogenic effects on hNCS in vitro, which we have also described here. Collectively, these findings demonstrate a pharmacological method for recapitulating improvements in BPS seen with other means of increasing AHN (20,25).

Immature adult-born granule cells from AHN have been shown to play a role in encoding contextual information (26,71,87) and in supporting distinct neural patterns for different contexts (88). While there may be acute effects of RO6871135 on behavior, these were not specifically tested here, and the treatment schedule was in fact designed to minimize these effects: RO6871135 was administered after behavioral testing and has a half-life of less than 4 hours. It is conceivable that acute or chronic RO6871135 treatment may have effects outside of the DG and increased AHN. However, we show that AHN effects are necessary for enhanced BPS, since RO6871135 does not improve BPS after ablation of AHN with targeted hippocampal irradiation. In these studies, much of the context discrimination is driven by increased freezing in the shock context, as mice continue to experience a foot shock on subsequent days of exposure. Increased locomotion with RO6871135 treatment refutes the explanation that these mice have a general increase in immobility. Likewise, decreased latency in the NSF test indicates that RO6871135 decreases anxiety-like behavior, and would not be expected to independently increase anxiety-related freezing. And while contextual fear conditioning after a single foot shock is unchanged with RO6871135, we demonstrate that the more challenging task of discriminating from a very similar context is where the neurogenic effects are most apparent, consistent with previous findings (25).

In vivo testing with 14 days of drug treatment significantly elevated Ki67 (a measure of proliferation) and BrdU (here indicating adult-born cell survival) cell count in the DG at two doses of the drug. DCX was only modestly elevated with the 14-day time course, and only at a higher dose. Given the known time course for DCX expression and maturation of adult-born neurons (89), DCX levels would be expected to peak or plateau after 3-4 weeks of treatment, which we observed in subsequent studies.

Aged mice show a deficit in BPS compared to young adult mice,(20,44–46) with unchanged response to foot shock (46), and they also have dramatically decreased levels of AHN (18,40–43). In humans, BPS deteriorates with age, and BPS impairments are correlated with cognitive impairment and dementia symptoms (50,55,63–66). In the present study, treatment with RO6871135 improved but did not restore neurogenesis levels or BPS performance to the same levels as young mice. However, these findings suggest that enhancing AHN may be an effective strategy to slow or reverse some of the cognitive effects of aging, and that neurogenic treatment can still be effective even in the setting of aging.

Pharmacologically-induced increases in AHN have been observed with serotonin reuptake inhibitors such as fluoxetine in the past, which were dependent partially on 5HT1A receptors expressed in the DG (69,90). Other antidepressant treatments such as tricyclic antidepressants (69,91) and monoamine oxidase inhibitors (92) also increase AHN. RO6871135 appears to have a novel mechanism of action, since the binding profile of RO6871135 showed no direct activity on the serotonin system except at very high concentrations. Indeed, treatment with antidepressants with no known serotonergic activity, such as norepinephrine-selective reuptake inhibitors and electroconvulsive stimulation have also demonstrated enhanced AHN (92). Some behavioral effects of SSRIs are also mediated through neurogenesis-independent mechanisms (72,90), and we show that RO6871135 effects in the NSF test were resistant to irradiation. We found that RO6871135 altered innate anxiety-like behavior more broadly in mice exposed to chronic corticosterone. This was consistent with our prior experience with genetically-enhanced AHN, which is able to reverse effects of chronic stress on anxiety-like measures, but does not alter these behaviors in unstressed mice (25,38). Enhanced neurogenesis has been shown to increase resilience to chronic stress (35–39), therefore the ability of novel neurogenic compounds to enhance resilience to other models of chronic stress should be tested in future studies.

As with many therapeutics, including many currently available medications, the exact targets and mechanisms of action for RO6871135 are not yet known. Looking at the kinases that were inhibited by RO6871135 in vitro or in situ, or those that bound specifically to the neurogenic analog, the strongest convergence was on cyclin-dependent kinases CDK8 and CDK11. CDKs and CDK inhibitors are instrumental in neural development, regulating cell fate and differentiation (93). There is evidence that CDK inhibition may be a promising target to upregulate AHN (94). CAMK2a and CAMK2b, two highly abundant proteins in the brain, are known to be crucial for learning and plasticity in mice (95–100), and normal neural development in humans (101–104). MAP2K6, also binding significantly to RO6871135 in this in situ assay, activates mitogen-activated protein kinase p38 (105) within the MAPK/ERK/JNK signaling cascades. Within these cascades, p38 has been shown to have a role in stress response, development, apoptosis, and senescence (106), and may even mediate age-related decline in AHN (107), though there has been some conflicting reports of the directionality of its effects (107–111). GSK3B binding was also seen in situ, and this kinase is involved in the Wnt/β-catenin pathway, a regulator of AHN (83,112,113) that may also be a promising target for counteracting neural loss in neurodegenerative disorders (114,115). Future studies will be needed to determine the precise mechanism of action and whether it is via single or multiple targets.

RO6871135 is not being developed as a clinical molecule in part because there was evidence of proliferation in mouse and rat livers in vivo (data not shown). Further preclinical screening tests would need to be pursued to derisk the safety profile of RO6871135, or chemical optimization could potentially increase central efficacy. However, the current study serves as a proof-of-concept for high-throughput screening and preclinical testing of neurogenic compounds for BPS effects.

Since an initial report of AHN in humans in 1998 (15), techniques for demonstrating evidence of AHN have continued to evolve (42,116–127). While the number of new cells might be quite low in human adults, (123), their impact on hippocampal circuitry may proportionally increase when the system is ‘challenged’, i.e. in the settings of stress, neurodegeneration, or other pathology (128–132). There is also evidence that, though proliferation of adult-born cells decreases with age, maturation time also lengthens, such that the total number of immature neurons in the system is still a significant proportion of cells (133,134). Moreover, AHN in mice may be a useful readout for interventions that affect broader hippocampal functioning in humans, with many neurogenic manipulations in mice (exercise, enrichment, antidepressants) having therapeutic or resilience-building properties in humans.

In this study, we demonstrate that pharmacologically enhancing AHN is a means for improving BPS. Studies of neural functioning during BPS in humans find that a homologous neural circuit is engaged (47–49,55), and investigations in clinical populations implicates this cognitive process as a promising therapeutic target (14,55–57,61,63–66,135–137). It has even been shown that perceived clinical response to antidepressants is correlated with improvements in BPS performance (138). While medications like SSRIs and other classes of antidepressants can increase AHN (69,91,92), they also have side effects that are not tolerable to many patients (139–141). Identification of compounds with novel neurogenic mechanisms may provide a means to increase AHN with a higher efficiency and reduced side effect profiles, ultimately increasing the effectiveness in individuals with insufficient response to existing medications. They may also provide direct and symptomatic treatment for individuals with BPS deficits that work in concert with other types of therapy, including pharmacotherapy, psychotherapy, neural modulation, and/or cognitive rehabilitation. Further translation of this treatment target should be pursued in clinical trials of neurogenic agents to directly test for improvements in BPS and assess how this correlates with general clinical improvement.

## Acknowledgements and Disclosures

This work was funded and supported by the National Institutes of Health (Grant Nos. K08MH122893 [Principal Investigator (PI): WC], R01MH068542 [PI: RH], RF1AG080818 [Co-PI: RH]) Funding and support also provided by the Hope for Depression Research Foundation [PI: RH, Co-Investigator WC].

We thank the Roche Neurogenesis team for their effort in the identification and characterizations of RO687715. The following authors have been employed at F. Hoffmann-La Roche AG while working on this project: MS, JW, AA, HF, SG, JL, DR, MG, SZ, LS, RJ.

Funding support was provided by Roche AG to BS, DJD, IMD, and RH to perform experiments. Compound RO6871135 was provided free of charge to WC, KT, BS, DJD, IMD, and RH from Roche AG. DJD reports having received lecture fees from Roche AG.

The authors report no potential conflicts of interest.

