## Supplementary Information for "Pharmacological Enhancement of Adult Hippocampal Neurogenesis Improves Behavioral Pattern Separation in Young and Aged Mice"

Chang *et al.*

### Supplementary Methods

**High throughput content in vitro screening of human neurogenesis**: Human neural stem/progenitor cells (hNSCs) were derived from human embryonic stem cells (hESCs) according to previously reported procedures (1,2). The well-characterized SA001 hESC line (NIH Registration Number 0085; Cellectis, Sweden)(3) was used to generate a large hNSC cell bank that was used the the high throughput screen (HTS). NSCs were generated from hESCs using a modification of the dual SMAD inhibition protocol (1,4). hESCs were dissociated and plated into AggreWell-800 plates (STEMCELL Technologies) at a density of 5,000 cells per microwell in a neuronal medium supplemented (NEP) with 10 mM Y-27632. After five days, aggregates were recovered, plated on polyornithine-/laminin-coated (PL) dishes in NEP medium and cultured for an additional three days to form neural rosettes. Rosettes were manually isolated and subsequently plated on PL dishes in NEP medium. Upon reaching confluence, cells were dissociated with 0.05% trypsin/EDTA solution (LifeTechnologies) and plated at 100,000 cells/cm$^2$ on PL dishes in medium containing FGF2, EGF and BDNF. Cells were cultured under these conditions for several passages. Medium was replaced daily throughout the entire neural progenitor cell (NPC) derivation procedure. NSC lines were characterized by FACS and immunocytochemical analyses including Nestin/Sox2 staining.

For the HTS, NSCs were thawed for 3 passages. Then the cells were seeded in Polyornithin/ Laminin coated 384 well plates at a cell density of 21,000 cells/cm$^2$ and a volume of 38µl. 4 hours after cell seeding, the compounds are added. Final DMSO concentration is 0.25%. The Assay Volume is 40µl/well. Controls were used including 100ng/ml FGF2, 20ng/ml EGF and 100ng/ml Wnt3a + 0.1%DMSO. After 4 days incubation at 37°C, 5% $CO_2$ the amount of ATP per well is quantified. The ATP concentration is proportional to the cell number. For the ATP quantification a commercial kit from Promega is used: CellTiterGlo®.

After the HTS, a hit confirmation was performed in a dose response (11 points, dilution factor is 2) and ranges from 8µM to 8nM and neurogenic specificity was interrogated testing on hNSCs with a counterscreen on mesenchymal stem cells (hMSCs). Subsequently, a High-Content-Screening (HCS) cellular imaging assay was used to detect cell number and immature neuronal differentiation with a stain for beta-tubulin and DAPI. Acquisition and automated quantification performed with Operetta HCS.

From this in vitro screening cascade, hits were then chemically modified and optimized based on multiple parameters including potency, safety assays and DMPK, which led to RO6871135. RO6871135 was tested for hepatic enzyme interactions, hERG toxicity, the micronucleus test and Ames tests for genotoxicity, a glutathione consumption assay. RO6871135 was pharmacokinetically profiled and selected for in vivo testing.

**Mice**: For the initial in vivo neurogenesis study, male 129/Sv mice were obtained from Taconic Farms (Denmark) at 8-10 weeks of age. For behavioral experiments, male c57BL/6J were obtained from Jackson Laboratories or Taconic at 8 weeks of age. For aging experiments, male c57BL/6J mice were obtained from the National Institute of Aging at 18 months of age and acclimated to the facility for at least 1 day after arrival. For chronic corticosterone experiments, male c57BL/6 NTac mice were purchased from Taconic Farms at 8-10 weeks of age. All mice were acclimated for at least 1 day after arrival. Mice were housed two to five per cage and maintained on a 12 h light/dark schedule with access to food and water *ad libitum*. All gavage and behavioral testing was conducted during the light cycle. Animal protocols for in vivo behavioral experiments without corticosterone were approved by the Institutional Animal Care and Use Committee at the New York State Psychiatric Institute and were in compliance with the NIH Guide for the Care and Use of Laboratory Animals. Chronic corticosterone experiments were conducted in compliance with protocols approved by another Institutional Animal Care and Use Committee (Council directive no. 87–848, 19 October 1987, Ministère de l’Agriculture et de la Forêt, Service Vétérinaire de la Santé et de la Protection Animale, permissions no. 92–256B to D. J. David). Care was taken to minimize both the number of animals used and their suffering.

**Drug administration**: RO6871135 was suspended in Klucel, 2% / Tween 80, 0.1% at a dose volume of 1.875 mg/ml. Aliquots of frozen compound were thawed and resuspended daily prior to oral gavage. For all behavioral experiments, 7.5 mg/kg of RO6871135 was administered daily. Histology studies in aged mice also included a 2.5 mg/kg group. After chronic treatment (21 days), the mice completed behavioral testing in the following order: Open Field, Novelty Suppressed Feeding, and Context Discrimination. On the days of behavioral testing, animals were gavaged after the cessation of the behavioral testing for the day to avoid any possible acute effects of drug treatment.

**In vivo neurogenesis study**: In young mice, vehicle or RO6871135 7.5 mg/kg or 15 mg/kg was administered by oral gavage for 14 days, with IP administration of BrdU 50 mg/kg for 5 days starting on Day 2. No behavioral testing was performed for this study Immunohistochemistry for these studies was performed to quantify BrdU (proliferation/survival), Ki67 (proliferation), and doublecortin (immature neurons). See below for immunohistochemistry methods.

**Open Field**: The mice were tested after 21 days after drug treatment. Before testing began, mice were allowed to habituate in their home cages in the testing room in the dark for 30 minutes. The mice were run in open field boxes from Kinder Scientific in the dark for 30 minutes and the data were collected and analyzed with MotorMonitor Software. Movement was captured via infrared detectors in the enclosures. Total distance traveled was analyzed for possible differences in locomotor activity among groups respectively.

For chronic corticosterone experiments, motor activity was quantified in Plexiglas open field boxes 43 x 43 $cm^2$ (MED associates, Georgia, VT) over a 30 min session. Two sets of 16 pulse-modulated infrared photo beams were placed on opposite walls 2.5 cm apart to record x-y ambulatory movements. A 40-W white bulb placed in the middle of the room provided around 200-lx illumination at floor level. Activity chambers were computer interfaced for data sampling at 100 ms resolution. The computer defined grid lines that divided each open field into center and surrounds regions, with each of four lines being 11 cm from each wall. Dependant measures were total time spent in the center, the numbers of entries into the center and distance travelled in the center divided by total distance travelled. Overall motor activity was quantified as the total distance travelled (cm).

**Novelty suppressed feeding**: Novelty Suppressed Feeding was performed as described (5,6). 24 hours prior to behavioral testing, all food was removed from the home cage. At the time of testing, a single food pellet attached to a food platform was placed in the center of the arena and a stopwatch was immediately started. The latency to eat (defined as the mouse sitting on its haunches and biting the pellet with the use of forepaws) was timed. The testing was stopped after 6 minutes, 10 minutes for corticosterone studies. Any animals that did not eat during the test were censored for survivial analysis. Immediately afterwards, the animal was transferred to its home cage, and the amount of food consumed by the mouse in the subsequent 5 min was measured. Each mouse was weighed before food deprivation and before testing to assess the percentage of body weight loss (data not shown).

**Contextual fear conditioning**: For young adult mice in the randomized fear discrimination paradigm, conditioning was conducted in a chamber from Coulborn Instruments (Model H10-24T) with internal dimensions of 7 x 7 x 12 in. The chambers had a clear plastic front and back walls, stainless steel walls on each side, and stainless-steel bars on the floor. For aged mice and young adult mice in the non-randomized paradigm, conditioning was conducted on one side of a Med-Associates shuttle box (ENV-010MC; 20.3cm X 15.9 cm X 21.3cm high) with a clear plexiglass wall, 3 aluminum walls and a stainless-steel grid as a floor. For all experiments, the chamber was encased by a sound-dampening cabinet. A house light was mounted directly above the chamber within the cabinet, and each chamber contained a ventilation fan. On the days of testing, mice were brought out of the vivarium and allowed to habituate in holding cages for at least 30 minutes before starting the experiment. Mouse behavior was recorded by digital video cameras mounted above the conditioning chamber. FreezeFrame and FreezeView software (Actimetrics) were used for recording and analyzing freezing behavior, respectively. Mice were allowed to freely explore the chamber for three minutes before delivery of a single 0.75 mA foot shock lasting 2 seconds. The mouse was taken out 15 sec after termination of the foot shock and returned to its home cage. The context for fear conditioning was designated as Context A. See *Tables S1 and S2 for parameters of all contexts. Fear conditioning was tested in the same Context A.

**Fear Generalization:** For young adult mice, a distinct Context C was used to assess fear generalization to a novel context. Compared to the fear-associated Context A, in Context C, the highly salient stainless-steel floor was covered with a plastic panel, the room lighting was much dimmer and illuminated with red lights, the mice were transported in a different shaped holding cage, the chamber light and ventilation fan were both off, a different scent was placed in the chamber, the cabinet door was left open, and a different cleaning solution was used to wipe the chamber between mice.

**Contextual Fear Discrimination**: The pattern separation task consisted of a contextual fear discrimination paradigm in which the mice had to learn to distinguish between a fearful shock context and a similar non-shock context, adapted from the methods described in (7). This was started the day after one-trial contextual fear conditioning was completed. Recall and distinct contexts were not tested in experiments where the fear discrimination contexts were not randomized, in order to make the task easier. See Tables S1 and S2 for parameters of all contexts. In pilot experiments, the similar context was found to evoke comparable levels of freezing behavior at that observed in the training context, suggesting generalization (pattern completion) between the two contexts. For discrimination learning, the timing, length, and strength of the foot shock was the same as for the initial learning day. The similar context B session was the same length, but with no foot shock delivered. Measurement of the freezing levels in both the training context (3-min pre-shock) and the similar context (3 min) each day allowed the assessment of discrimination between the two contexts.

**Immunohistochemistry of neurogenesis markers**: Mice were anesthetized with ketamine/xylazine (100 mg/ml ketamine, 20 mg/ml xylazine) and transcardially perfused with 10 ml cold saline, followed by 30 ml ice-cold paraformaldehyde (PFA; wt/vol) in saline. Brains were dissected and post-fixed overnight in 4% PFA at 4°C. Brains were then cryoprotected in 30% sucrose (wt/vol) for three days at 4°C and subsequently frozen in optimum cutting temperature (OCT) compound (Tissue Tek, Torrance, CA) and stored at -80 °C until cryostat sectioning. Serial sections (45 μM) were cut through the entire hippocampus on a cryostat (Leica CM3050S) and stored in phosphate-buffered saline (PBS) with 0.1% $NaN_3$ until further processing.

For the 14-day in vivo neurogenesis test and the young vs aged mouse studies, tissue and cells were fixed and processed for immunostaining as previously described (8). Primary antibodies and dilutions used for this study included goat anti-doublecortin (DCX) (1:200, Santa Cruz Biotechnology), rabbit anti-Ki67 (1:200, Vision BioSystems) and rat 1:400; rat anti-BrdU (Accurate). Secondary antibodies coupled to the fluorophores Cy3, Cy5, FITC, or Alexa 488 were obtained from The Jackson Laboratory and were used at a dilution of 1:250 after resuspension in 200 µl of $H_2O$ and 200 µl of glycerol. Quantification was obtained on the Leica SP5 confocal microscope.

For BrdU labeling studies in aged versus young adult mice, BrdU was administered once a day intraperitoneally in 0.9% NaCl for 5 days starting after 14 days of drug treatment, prior to behavioral testing. The same methods were used for immunohistochemical quantification of neurogenesis markers as above.

For doublecortin (DCX) staining in young adult mice after randomized fear discrimination task behavioral testing, antigen retrieval was performed prior to staining: sections were washed three times for 10 min in PBS and placed in sodium citrate buffer at pH 8 and heated to 100 °C, then incubated in 80 °C for 30 minutes, then allowed to return to room temperature. Sections were washed again three times for 10 min in PBS prior to incubation in 10% normal donkey serum (NDS, wt/vol) in 0.2% Triton X-100 (PBST) for 2 hours, then incubated overnight at 4 °C in primary antibody in PBST + 10% NDS (rabbit anti-doublecortin 1:600, Cell Signaling). The next day, sections were washed three times for 10 min in PBS and incubated in secondary antibody (1:400 anti-rabbit Alexa Fluor 488, Jackson ImmunoResearch) in PBST + 10% NDS for 2 hrs at room temperature. Sections were rinsed for 10 min in 1:1000 Hoechst in PBS (Invitrogen) to label cell nuclei, then mounted onto glass slides and coverslipped with ProLong Glass antifade reagent (Invitrogen). Histology slides were imaged on a (Leica TCS SP8) confocal microscope using a 10x or 20x objective.

For DCX staining in young adult mice after chronic corticosterone treatment, fixed and sliced sections were incubated in 0.1M TBS with 0.5% Triton X-100 and NDS for 1 hour, followed by anti-rat doublecortin primary antibody (1:100) in TBS/TritonX for 24 hours at 4°C. The secondary antibody was biotinylated donkey anti-goat (1:500) in TBS/NDS for 1 hr at room temperature, followed by a 1 hr amplification step using an avidin-biotin complex (Vector, USA). The immunohistochemistry protocol was adapted from David et al. (5).

**Histology analysis**: For the 14-day in vivo neurogenesis test and the young vs aged mouse studies, cells positive for BrdU, Ki67, and DCX were manually counted from 6-10 sections per mouse. For young mice that underwent behavioral testing, DCX staining was detected automatically using a threshold in ImageJ within a region of interest manually drawn around the DG granule cell and molecular layers. DCX-positive ($DCX^+$) pixels were divided by the total area of the ROI. Both hemispheres of 4-6 sections from the dorsal and ventral DG. For chronic corticosterone experiments, $DCX^+$ cells were manually counted. Six sections per animal were analysed: 2 sections in the dorsal hippocampus, 2 sections in the ventral hippocampus and 2 sections in the middle of the structure. All quantification was performed blind to group identity. Values from each mouse were averaged prior to statistical analyses.

**Chronic corticosterone paradigm:** Corticosterone was administered in drinking water (35 µg/ml in 0.45% β-cyclodextrine drinking water) with controls receiving 0.45% β-cyclodextrine (β-CD) vehicle. After 4 weeks, mice were co-treated with corticosterone or β-CD and either RO6871135 7.5 mg/kg or the vehicle by oral gavage until the completion of the study. Behavioral testing began 4 weeks after starting drug treatment.

**Focal X-ray irradiation of the hippocampus**: 8-week old mice were anesthetized with sodium pentobarbital (administered intraperitoneally at 0.6 mg/ml concentration, 6 mg/kg body weight once a day for three days, each injection spaced apart by 3 days), placed in a stereotaxic frame and exposed to cranial irradiation using a Precision X-Ray X-RAD320 system. There was a 3.22 x 11 mm window above the hippocampus in a lead plate that otherwise shielded the entire body and allowed for focal X-ray applications. X-rays were filtered using a 2 mm Al filter, the corrected dose rate was approximately 1.8 Gy per min and the source to skin distance was 30 cm. A cumulative dose of 5 Gy was given over the course of 2 minutes and 47 seconds on days 1, 4, and 8. Drug treatment was initiated 8 weeks after hippocampal x-irradiation, and behavioral testing was carried out 3 weeks after that.

**In vitro off-target screening assay:** In vitro pharmacological screening for off-target assays were also performed as described (9), tested at Cerep (now Eurofins Pharma Discovery). These assays test for potential hazards of a compound and tests G protein-coupled receptors, ion channels, nuclear hormone receptors, transporters, and some enzymes correlated with off-target related adverse drug reactions (10). Single dose percentage inhibition for 10 µM test concentration of RO6871135 were assessed in duplicate using radioligand binding displacement and enzyme assays, with ≥ 50% inhibition deemed to show activity at that off-target cite. The experiment was accepted in accordance with Eurofins validation standard operating procedure. The $IC_{50}$ values (concentration causing a half-maximal inhibition of control specific binding) and Hill coefficients (nH) were determined by non-linear regression analysis of the competition curves generated with mean replicate values using Hill equation curve fitting.

**In vitro kinase screening assay:** Kinase activity inhibition by RO6871135 at 1 μM was profiled against 96 human kinases as described (11) via LeadHunter® Drug Discovery Services Panels (Eurofins DiscoverX Products, LLC, Fremont, CA). Briefly, a solid support competitively binds kinase against the test compound, RO6871135. The amount of kinase captured in test versus control samples is measured by qPCR that detects for an associated DNA label. Dissociation constants ($K_d$) for compound-kinase interactions are calculated by measuring the amount of kinase captured on the solid support as a function of the test compound concentration. CDK8 was added for $K_d$ determination due to special interest based on other binding and activity assays.

**In situ kinase binding assay:** Kinome-wide binding for possible mechanisms of action was conducted using the KiNativ™ platform (12–14), performed by ActivX. Brain and liver tissue samples were collected from mice 4 hours after treatment with vehicle or 7.5 mg/kg RO6871135 (*N*=4 mice/group). 0.5 mL of tissue lysate from each sample at ~5mg/ml was labeled by the addition of 5 μM probe and incubated for 15 minutes. Proteins are quickly denatured with Urea/DTT and processed for proteolytic digestion. Following a digestion with Trypsin, the samples are affinity captured to isolate all probe-bound peptides. Peptides are then analyzed using customized liquid chromatography–mass spectrometry (LC-MS/MS) protocols designed to target kinase-peptide-probe conjugates that are known to be present. Signals from these kinase peptides are quantified by extracting the ion traces of multiple predicted fragment ions and integrating the resulting peak areas. ATP probes were used and analyzed with the mouse brain-ATP and mouse liver-ATP target listings. As previously described (12), a cutoff of 35% inhibition, plus a significance value of ≥0.8 was used, which had been found to yield <1% false positive rate.

**Chemical proteomics analysis**: Human neuronal stem cells (hNSC) differentiated for 80 hours were cultivated in a 1:1 DMEM/F12/Neurobasal medium containing B27 supplement without Vitamin A, N2 supplement, beta-mercaptoethanol, FGF, EGF and BDNF,and either light (unlabeled) L-Lysine and L-Arginine, or heavy L-Lysine 8 (13C6 ; 15N2) and Heavy Arginine 10 (13C6; 15N4), on Poly-L-Ornithine/Laminin coated flasks. Modified piperazinone compounds (active RO6932557 and inactive RO6954205) were coupled to NHS Sepharose 4B according to the manufacturer’s instructions. Cell pellets were resuspended in lysis buffer (Hepes 50mM NaCl 150 mM CaCl2 1mM MgCl2 1mM 0.5%NP40 Complete EDTA free and PhosphoSTOP),

20 min on ice. Lysates were cleared by centrifugation before loading onto the Sepharose beads. The lysate-beads suspension was left at 4°C under agitation for 1 hour. After 3 washes with lysis buffer, proteins were eluted from the beads at 85°C in reducing SDS-PAGE loading buffer. Respective Heavy and Light elutions were pooled and elution fractions were subsequently separated by SDS-PAGE. Each gel lane was cut into 4 bands which were reduced in dithiothreitol, alkylated with iodoacetamide and digested with trypsin. Peptides were extracted with ammonium bicarbonate 25mM and formic acid 5%. After drying, peptides were resuspended in 2% acetonitrile, 5% formic acid and analyzed in two technical replicates by LC-MS/MS using a nanoflow liquid chromatography system (Proxeon EASY-nLC) coupled to an electrospray ion trap mass spectrometer (Velos LTQ-Orbitrap). Raw data files were processed using Mascot (version 2.4). Statistical analysis for chemical proteomics was conducted using R 4.3.2, R: A language and environment for statistical computing. R Foundation for Statistical Computing, Vienna, Austria, and packages 'XML' (version 3.99), 'plyr' (version 1.8.9) and 'lattice' (version 0.21). To avoid redundancy over bands, for each protein the most abundant band was selected for analysis. For samples with reversed H/L, the ratios were harmonized by 1/(L/H) = H/L. H/L values were $\log_2$-transformed, then centered by the respective median and standardized by the median absolute deviation (mad). For each protein, a one-sample t-Test was computed, with the null hypothesis $\mu = \log_2(1{:}1) = 0$ and alternative hypothesis $\mu > \log_2(1{:}1) = 0$. p-Values were adjusted following Benjamini, Y., and Yekutieli, D. (2001, The control of the false discovery rate in multiple testing under dependency. Annals of Statistics, 29, 1165–1188).

**Statistical Analysis**: In addition to statistical analyses described above, additional analyses were carried out using the Python packages statsmodels (15) and SciPy (16) and in GraphPad Prism (version 9.5.1 for macOS, GraphPad Software, San Diego, California, USA, www.graphpad.com.) Statistical significance was assessed for histology measures by repeated measures analysis of variance (ANOVA) tests followed by Dunnett's multiple comparisons test. Statistical significance was assessed in the fear discrimination task by repeated-measures two-way analysis of variance ANOVA, followed by Šidák's multiple comparison test for individual days. Statistical significance for single-day tasks were assessed by unpaired two-tailed student's t-tests or analysis of variance (ANOVA). Significant main effects or interactions were followed up with Šidák's multiple comparison test or Tukey's method for *post hoc* tests, where appropriate. Statistical significance for the NSF task was assessed using Log-rank (Mantel-Cox) test, and when multiple comparisons were used, Bonferroni correction of $\alpha$ was applied.

Behavioral testing, scoring, and analysis were performed blind to treatment as much as possible.

### Supplementary Figures

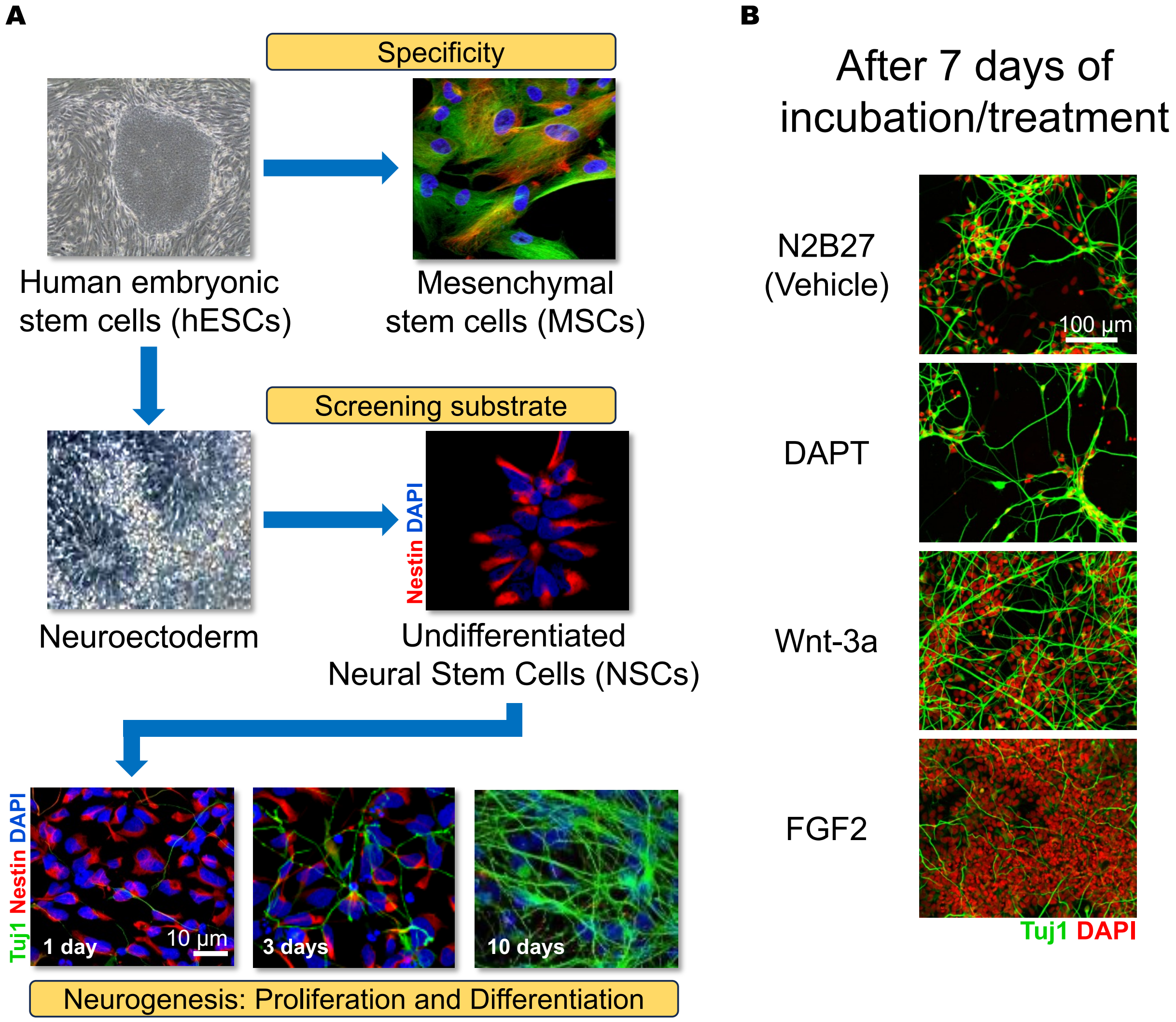


**Figure S1:** In vitro human embryonic stem cell (hESC) derived neural stem cells (NSCs) as a cell model to study human neurogenesis. (A) Outline of the protocol for derivation of neural stem cells from hESC via dual smad differentiation protocol as previously described (1) with representative microscopic and confocal images of each stage. Undifferentiated NSCs are

Nestin-positive (red). In the bottom panels, NSCs are Nestin-positive (red) and Tuj1-negative (green), while 3-10 days after withdrawal of growth factors there is a loss of Nestin staining concomitant with an upregulation of the expression of Tuj1, indicating neuronal differentiation. (B) NSCs respond to neurogenic pathway modulators. Representative confocal images of 7-day-old differentiated neurons cultured under the labeled conditions to show responsiveness of NSCs to known neurogenesis modulators. Control conditions shown for comparison. DAPT = N-[N-(3,5-difluorophenacetyl)-L-alanyl]-*S*-phenyl-glycine t-butyle ester, a γ-secretase inhibitor that blocks Notch signaling and accelerates differentiation (17). Wnt-3a promotes proliferation and neuronal differentiation (8,18). FGF2 = fibroblast growth factor-2, which stimulates proliferation.

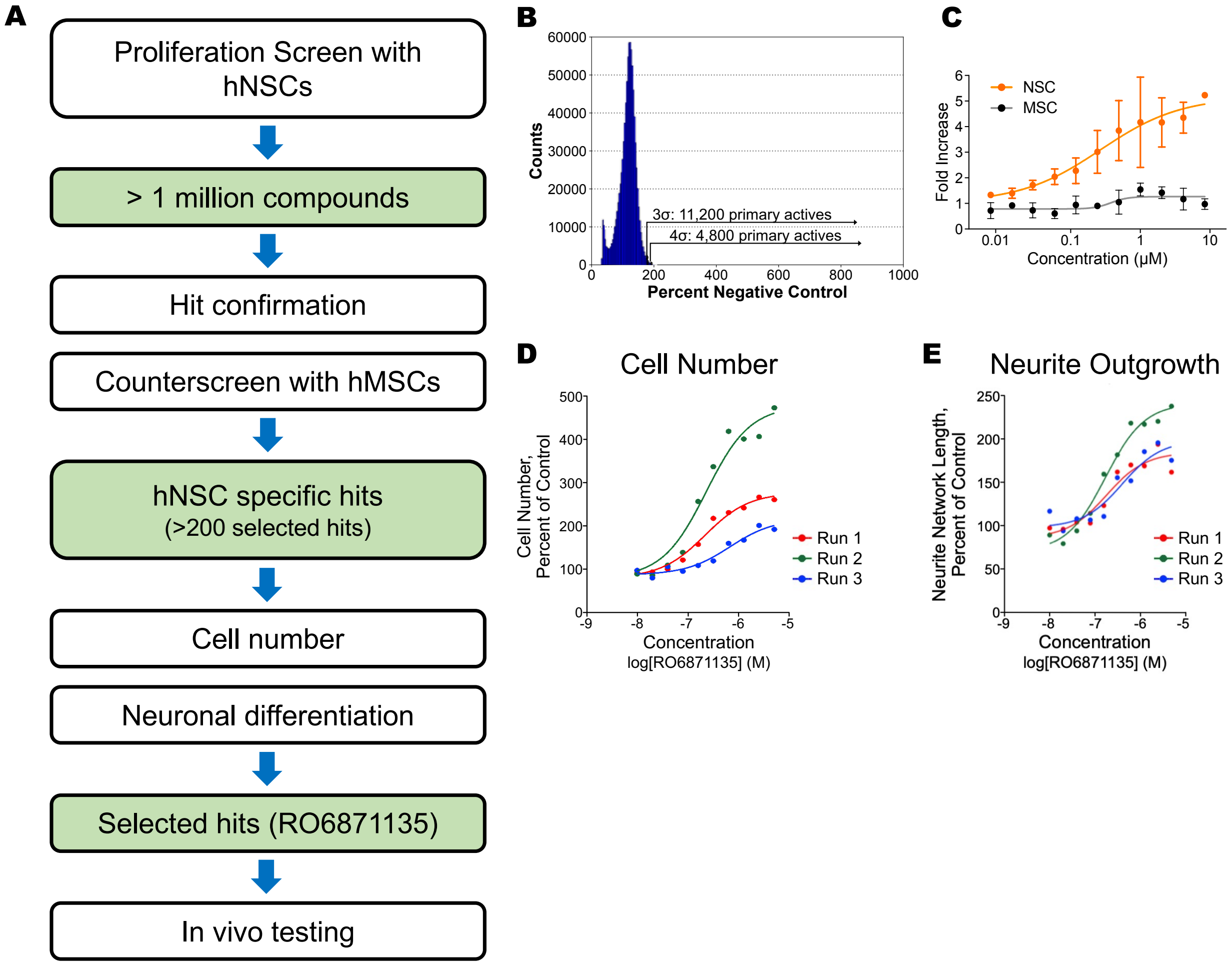

**Figure S2:** Screening cascade to identify novel for neurogenic compounds. (A) Screening work flow. (B) Results of high-throughput screen with hNSCs. Positive hits depicted from 3 or 4 standard deviations from the mean. (C) Dose-response confirmation of effects of RO6871135 in hNSCs over hMSCs demonstrating neural selectivity. Shown is the average of two repetitions and best fit curves. (D) Testing of dose-response for RO6871135 in high-content screen for cell number of hNSCs. Cell number values calculated from nuclei density (%area of well), normalized to DMSO control values, which were set to 100%. (E) Testing of dose-response for RO6871135 increasing neuronal differentiation, as measured by total neurite network length. Neurite network lengths were normalized to DMSO control values, set to 100%. hNSC = human embryonic stem cell derived neural stem cells; hMSC = human embryonic stem cell derived mesenchymal stem cells.

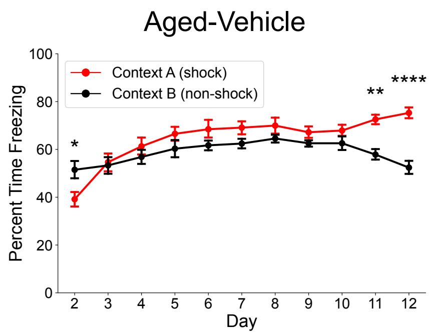


**Figure S3:** Aged vehicle-treated mice eventually learn to discriminate between contexts A and B. With additional days of the fear discrimination task, aged vehicle-treated mice still have a significant context × day interaction ($F_{(10,462)}$= 10.39, $p$<0.0001). Post hoc tests indicated they significantly distinguished between the similar contexts, as measured by percent time freezing, on days 11 and 12 . *$p$<0.05, **$p$<0.01, ***$p$<0.001, ****$p$<0.0001.

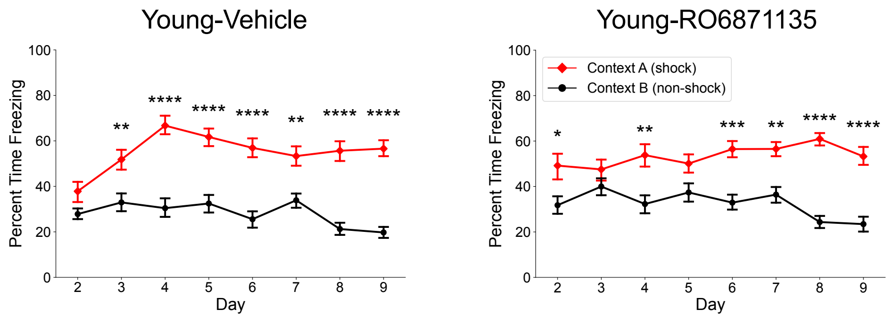


**Figure S4:** In the non-randomized fear discrimination paradigm, RO6871135 did not exhibit accelerated pattern separation in young adult mice. Data from Figure 4 vehicle-treated young mice shown again here for reference. Young mice treated with RO6871135 also had a significant context × day interaction ($F_{(7,196)}$= 4.435, $p$<0.0001), and displayed higher freezing in Context A on Day 2. However, this difference was not maintained consistently until Day 6. Discrimination between contexts in vehicle- vs. RO6871135-treated mice was not qualitatively different between groups, perhaps due to near-maximal performance in the vehicle-treated group in this easier, non-randomized version of the task. *$p$<0.05, **$p$<0.01, ***$p$<0.001, ****$p$<0.0001.

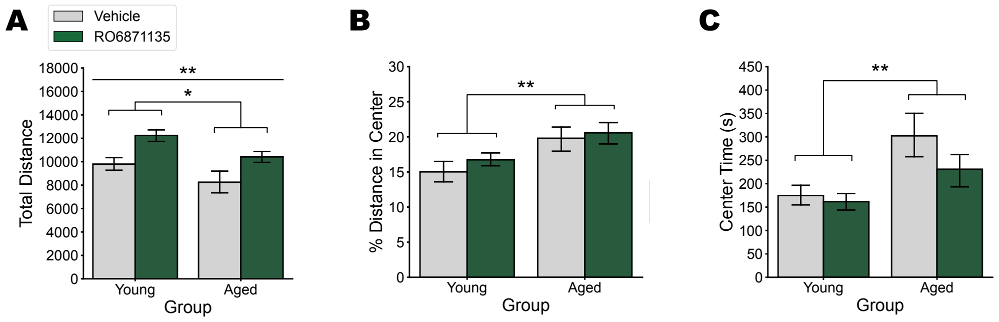


**Figure S5:** Locomotion is increased by chronic RO6871135, while innate anxiety the Open Field Test (OFT) is unchanged. (Left Panel): There was an overall effect of age ($F_{(1,54)}$= 6.182, $p$<0.05) and drug treatment ($F_{(1,54)}$= 11.518, $p$<0.005) on total distance travelled in the open field after treatment with RO6871135, but no significant interaction of age × drug ($F_{(1,54)}$= 0.044, NS). (Center Panel): Aged mice had increased percent distance in OFT center ($F_{(1,54)}$= 8.538,

$p$<0.01). (Right Panel): Aged mice had increased time spent in the center of the OFT ($F_{(1,54)}$= 9.32, $p$<0.005). There was no effect of drug group on distance or time in the center (%distance $F_{(1,54)}$= 1.566, NS; center time $F_{(1,54)}$= 0.772, NS), and no age × drug interaction ( %distance $F_{(1,54)}$= 0.744, NS; center time $F_{(1,54)}$= 0.111, NS). *$p$<0.05, **$p$<0.01.

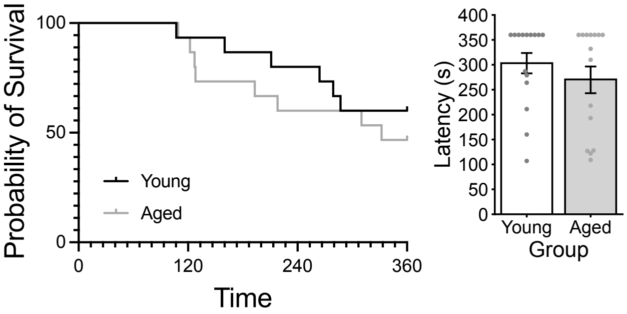


**Figure S6:** Aging does not alter performance in the NSF test. Young adult and aged mice show similar levels of latency to feed in the arena, represented as a survival curve. Log-rank (Mantel-Cox) test: Chi square = 0.5875, *NS*. Latency values for individual mice are also shown on the bar graph. This is a direct comparison of vehicle-treated mice also shown in Figure 3F, G.

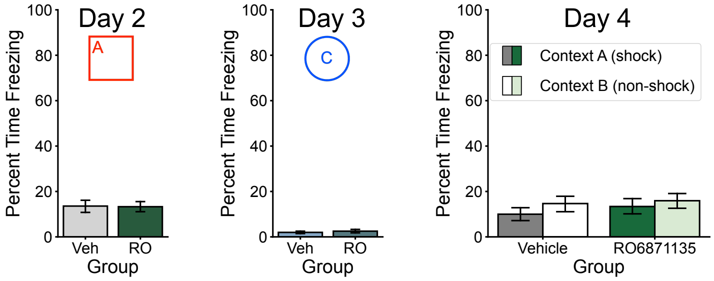


**Figure S7:** Irradiation does not alter contextual fear conditioning ($t$(15)= 0.079, NS) or increase generalization of freezing in a different, novel context ($t$(15)= -0.601, NS). Both groups demonstrate similar levels of freezing on Day 4 in Contexts A and B (context $F_{(1,30)}$= 1.083, NS; group $F_{(1,30)}$= 0.446, NS; group × context $F_{(1,30)}$= 0.095, NS).

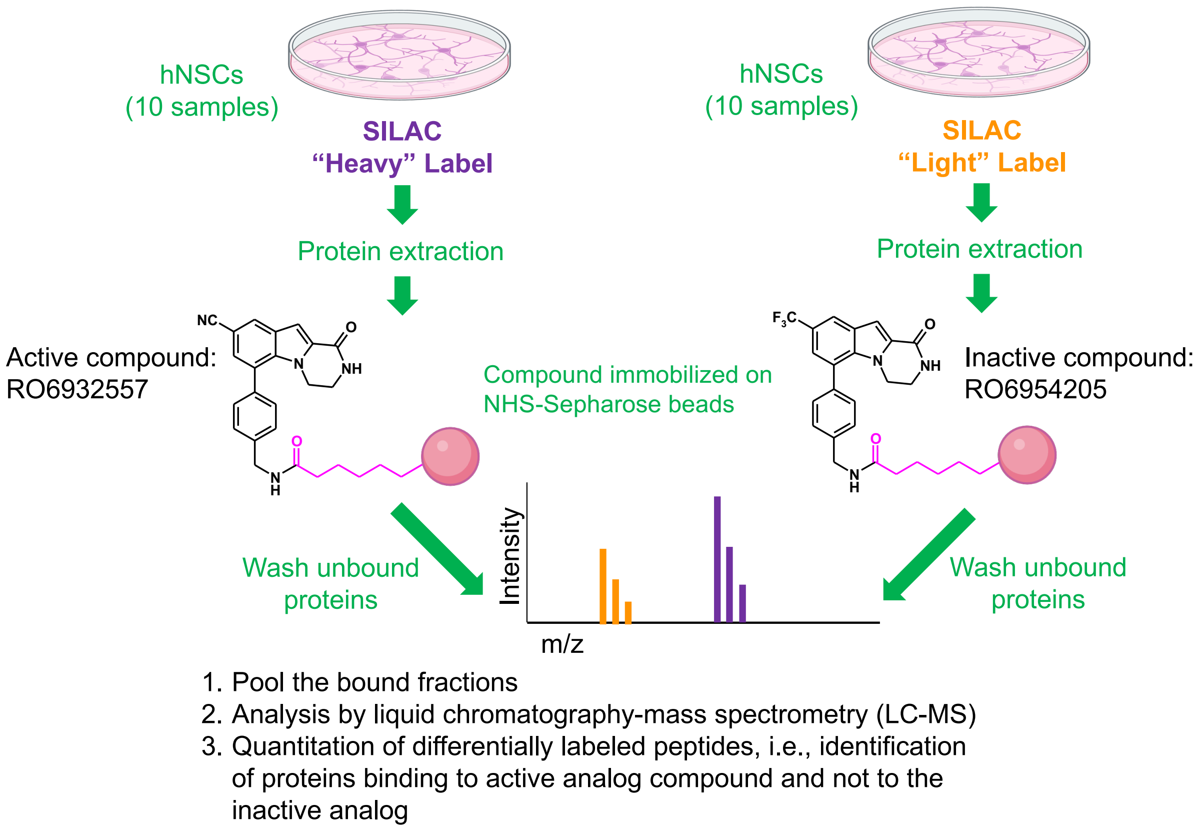


**Figure S8:** Chemical proteomics study for determining the in vitro human NSC binding profile of an active piperazinone analog to RO6871135. Two modified versions of RO6871135, RO6932557 (active for neurogenesis in vitro) and RO6954205 (inactive analog), were synthesized and covalently bound to NHS-Sepharose beads via short linker (shown in magenta in the molecular structure). Human embryonic stem cell derived neural stem cells (hNSCs) were differentially labeled with stable isotope labeling using amino acids in cell culture (SILAC) (19). Differentially labeled cell lysates were loaded either on the active or on the inactive compound matrix. Bound fractions were pooled and analyzed by mass spectrometry. A differential abundance analysis defined the proteins significantly enriched on the active compound (heavy SILAC label, purple) versus the inactive one (light SILAC label, orange).

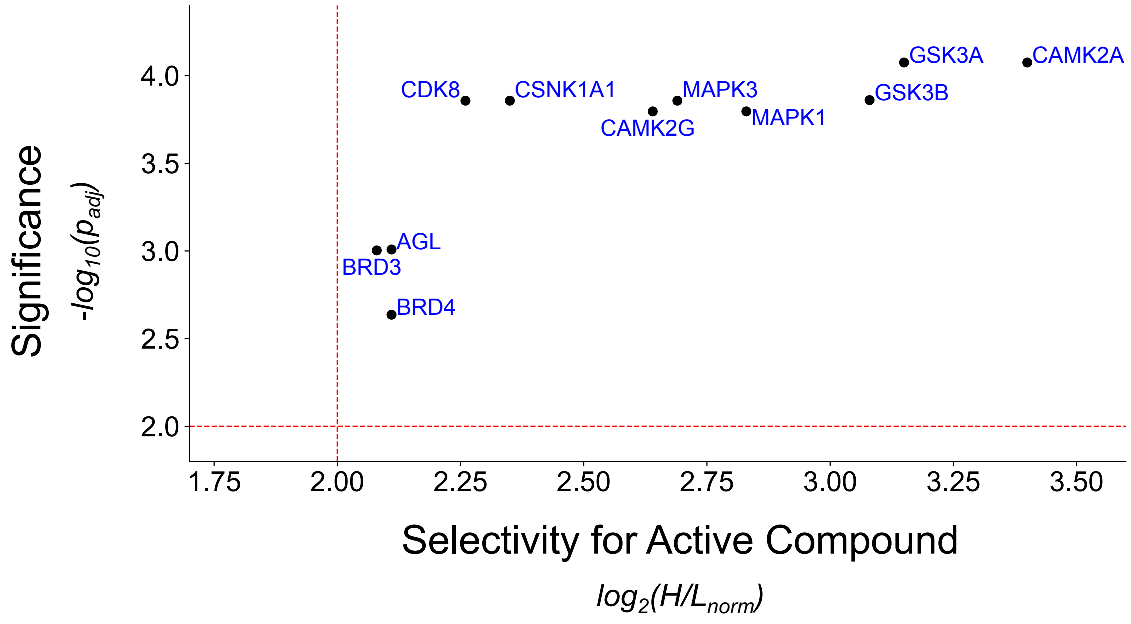


**Figure S9:** The protein binding profile of an active piperazinone (heavy labeled hNSC) analog over the inactive analog (light labeled hNSC) using a chemical proteomics method. The ratio of heavy labeled proteins (bound to the active compound) versus light labeled proteins (bound to the inactive compound) is shown as $log_2(H/L_{norm})$ on the x-axis. The red dashed line indicates a 4-fold binding difference for the active analog. Statistical significance, plotted as $-log_{10}(p_{adj})$. Points above the y-axis red dashed line have $p<0.05$. Proteins with statistically significant, high levels of binding in the neurogenic analog to RO6871135 indicate possible mechanisms of action RO6871135 and targets for additional neurogenic compounds.

### Supplementary tables

**Table S1:** Parameters for fear discrimination contexts in the random order paradigm, used for studies of young adult mice. The conditions for Contexts A (shock), B (non-shock, similar), and C (non-shock, novel) are listed.

| Randomized Order Fear Discrimination Task | | | |
|---|---|---|---|
| | **Context A** | **Context B** | **Context C** |
| **Room Light** | On | On | Red |
| **Chamber Light** | On | Off | Off |
| **Chamber Fan** | On | Off | Off |
| **Odor** | Satusuma | Strawberry | Coconut |
| **Floor** | Uncovered | Uncovered | Covered |
| **Chamber door** | Closed | Open | Open |
| **Chamber walls** | Uncovered | Covered | Uncovered |
| **Cleaner** | Virkon | Virkon | Sani-Cloths |
| **Holding cage** | Rectangular | Rectangular | Pie cages |

**Table S2:** Parameters for fear discrimination contexts in the non-randomized order paradigm, used for studies of aged mice. The conditions for Contexts A (shock) and B (non-shock, similar) are listed.

| Non-randomized Order Fear Discrimination Task | | |
|---|---|---|
| | **Context A** | **Context B** |
| **Plexiglas Wall** | Up | Down |
| **Chamber Light** | On | Off |
| **Chamber Fan** | On | Off |
| **Odor** | Anise | Lemon |
| **Floor** | Uncovered | Uncovered |
| **Chamber door** | Closed | Open |
| **Chamber walls** | Uncovered | Covered |
| **Cleaner** | Sani-Cloths | Ethanol |
| **Holding cage** | Rectangular plastic | Paper bucket |

**Table S3:** Additional profiling of RO6871135, including molecular properties, additional in vitro screens, and in vivo pharmacokinetics in mice. RO6871135 was tested for hepatic enzyme interactions, hERG toxicity, the micronucleus test and Ames tests for genotoxicity, a glutathione consumption assay.

<table>
<tr>
<td rowspan="2">Molecular properties:<br>Molecular weight: 345.2<br>Lipophilicity (cLogP): 5.4<br>Solubility: Lysa: <1 µg/mL<br>Thesa: <1 µg/mL<br>Fasted solubility (FaSSIF): 309 µg/mL<br>Feasted solubility (FeSSIF): 2983 µg/mL<br>Permeability (PAMPA): medium-high<br>Maximum achievable bioavailability (MAB):<br>human microsomes: 97%<br>mouse microsomes: 78%<br>Bioavailability (Fh, mouse hepatocytes): 41%</td>
<td>Potency in in vitro screen:<br>ATP- hNSCs: 0.026/607 µM<br>hMSC: >8/130 µM</td>
</tr>
<tr>
<td>In vitro Safety<br>CYP (3A4/2D6/2C9):<br>$IC_{50}$ (µM) > 50/47/25<br>hERG: $IC_{20}$ > 1µM<br>MNT: Negative<br>GSH: Negative<br>Ames: Negative</td>
</tr>
<tr>
<td colspan="2">Single Dose Pharmacokinetics (mouse)<br>$T_{1/2}$ = 3.7 hours<br>Clearance = 7.3 ml/min/kg<br>Steady state volume of distribution (Vss) = 2.48 l/kg<br>Bioavailability (F) = 100% (10 mg/kg p.o.)<br>Fu = BLQ (below limit of quantitification)<br>Kp (brain/plasma) = 1.7<br>P – gp: ER (human/mouse) = 1/1</td>
</tr>
</table>

**Table S4:** Top nine results from the off-target in vitro screening panel, showing that activity required much higher concentrations of RO6871135 than what was measured in vivo (6.5 [nM] in brain). Agonist or antagonist activity from a selected panel of adverse drug effect targets (Eurofins/Cerep panel), with corresponding $IC_{50}$ and $K_i$ concentrations as well as Hill coefficients ($n_H$). Full list of assays listed in Bendels et al., (9).

| Assay | Radioligand Type | $IC_{50}$ (μM) | $K_i$ (μM) | $n_H$ |
|---|---|---|---|---|
| $A_{2A}$ | Agonist | 2.6 | 2.1 | 1.5 |
| $A_3$ | Agonist | 1.3 | 0.8 | 1.3 |
| CXCR2 (IL-8B) | Agonist | 6.2 | 2.9 | 2.5 |
| $M_3$ | Antagonist | 5.1 | 3.7 | 2.5 |
| 5-$HT_{2B}$ | Agonist | 1.4 | 0.7 | 1.0 |
| 5-$HT_{2C}$ | Antagonist | 9.9 | 3.3 | 0.9 |
| 5-$HT_{5A}$ | Agonist | 1.7 | 0.9 | 1.4 |
| Norepinephrine transporter | Antagonist | 1.6 | 1.2 | 1.2 |
| Dopamine transporter | Antagonist | 0.8 | 0.4 | 1.2 |

**Table S5:** Full list of kinases screened for binding to RO6871135 with the in vitro KINOMEscan™ platform.

| KINOMEscan™ Kinase Assays | | |
|---|---|---|
| ABL1(E255K)-phosphorylated | FAK | PCTK1 |
| ABL1(T315I)-phosphorylated | FGFR2 | PDGFRA |
| ABL1-phosphorylated | FGFR3 | PDGFRB |
| ACVR1B | FLT3 | PDPK1 |
| ADCK3 | GSK3B | PIK3C2B |
| AKT1 | IGF1R | PIK3CA |
| AKT2 | IKK-alpha 80 | PIK3CG |
| ALK | IKK-beta | PIM1 |
| AURKA | INSR | PIM2 |
| AURKB | JAK2(JH1domain-catalytic) | PIM3 |
| AXL | JAK3(JH1domain-catalytic) | PKAC-alpha |
| BMPR2 | JNK1 | PLK1 |
| BRAF | JNK2 | PLK3 |
| BRAF(V600E) | JNK3 | PLK4 |
| BTK | KIT | PRKCE |
| CDK11 | KIT(D816V) | RAF1 |
| CDK2 | KIT(V559D,T670I) | RET |
| CDK3 | LKB1 | RIOK2 |
| CDK7 | MAP3K4 | ROCK2 |
| CDK9 | MAPKAPK2 | RSK2(Kin.Dom.1-N-terminal) |
| CHEK1 | MARK3 | SNARK |
| CSF1R | MEK1 | SRC |
| CSNK1D | MEK2 | SRPK3 |
| CSNK1G2 | MET | TGFBR1 |
| DCAMKL1 | MKNK1 | TIE2 |
| DYRK1B | MKNK2 | TRKA |
| EGFR | MLK1 | TSSK1B |
| EGFR(L858R) | p38-alpha | TYK2(JH1domain-catalytic) |
| EPHA2 | p38-beta | ULK2 |
| ERBB2 | PAK1 | VEGFR2 |
| ERBB4 | PAK2 | YANK3 |
| ERK1 | PAK4 | ZAP70 |

**Table S6:** Top results from KINOMEscan screening platform for RO6871135. At 1μM, RO6871135 demonstrated 96% inhibition of CDK11 and <50% for all other kinases in the panel. $K_d$ values listed here. There were no additional interactions with additional kinases in the 96 kinase panel.

| Assay | $K_d$ (nM) |
|---|---|
| CDK11 | 94 |
| CDK8 | 220 |
| CDK2 | > 40,000 |
| CDK4 | > 40,000 |

| Kinase | Activity |
|---|---|
| CaMK1a | 28.5 |
| CaMK1d | 50.3 |
| **CaMK2a** | 60.9 |
| **CaMK2b** | 54.2 |
| CaMK2d | 25.1 |
| CaMK2d | 25.4 |
| CaMK2g | 50.6 |
| CaMK4 | 29.6 |
| **CDK11,CDK8** | 66.8 |
| CDK5 | 9.5 |
| CDK5 | 8.0 |
| CDK5 | 27.4 |
| GSK3A | 15.4 |
| GSK3B | 38.8 |
| MAP2K1,MAP2K2 | 14.0 |
| MAP2K2 | -5.0 |
| MAP2K4 | -1.3 |
| MAP2K4 | 40.0 |
| **MAP2K6** | 53.2 |
| MAP3K2,MAP3K3 | -17.2 |
| MAP3K5 | 20.3 |
| MAP3K5,MAP3K6 | 5.2 |
| FER | 43.9 |
| FER | 45.6 |
| FER | 46.4 |
| ROCK1 | 0.0 |
| ROCK1,ROCK2 | 41.6 |
| TLK1 | 46.6 |
| ULK3 | 37.5 |

| |
|---|
| >90% inhibition |
| 75 - 90% inhibition |
| 50 - 75% inhibition |
| 35 - 50% inhibition |
| No change |
| Variable or weak data* |

**Table S7:** Brain in situ KiNativ$^{TM}$ profiling assay to suggest possible mechanisms of action for RO6871135. Highlighted significant results from a kinome-wide binding profile for brain tissue from mice treated with RO6871135 compared to tissue from vehicle-treated mice are shown here, with some comparison kinases from the same family also included. KiNativ$^{TM}$ profiling assay was performed as described by Patricelli et al. (12). RO6871135 has significant binding with >50% inhibition for CDK8 and CDK11, as well as to CaMK2a, CaMK2b, and MAP2K6. For full results and kinase descriptions, see Table S6.

**Table S8:** Full KiNativ™ profiling results of brain and liver tissue from mice treated with RO6871135 *vs*. vehicle.

| Legend |
|---|
| >90% inhibition |
| 75-90 % inhibition |
| 50-75% inhibition |
| 35-50% inhibition |
| No change (<35 %) |
| Variable or weak data |

| Kinase | Description | Sequence | Brain | Liver |
|---|---|---|---|---|
| AKT1 | RAC-alpha serine/threonine-protein kinase | GTFGKVILVK | -12.8 | |
| AKT2,AKT3 | RAC-beta serine/threonine-protein kinase | GTFGKVILVR | -22.1 | |
| AMPKa1 | 5'-AMP-activated protein kinase catalytic subunit alpha-1 | DLKPENVLLDAHMNAK | -29.4 | 12.8 |
| AMPKa1,AMPKa2 | 5'-AMP-activated protein kinase catalytic subunit alpha-1 | VAVKILNR | -2.3 | |
| AMPKa2 | 5'-AMP-activated protein kinase catalytic subunit alpha-2 | DLKPENVLLDAQMNAK | 15.9 | |
| BRAF | B-Raf proto-oncogene serine/threonine-protein kinase | DLKSNNIFLHEDLTVK | 33.8 | -16.2 |
| BRSK2 | BR serine/threonine-protein kinase 2 | VAIKIVNR | 28.6 | |
| BRSK2 | BR serine/threonine-protein kinase 2 | DLKPENLLLDER | 2.4 | |
| CaMK1a | Calcium/calmodulin-dependent protein kinase type 1 | LVAIKCIAK | 28.5 | 2.3 |
| CaMK1d | Calcium/calmodulin-dependent protein kinase type 1D | LFAVKCIPK | 50.3 | 9.6 |
| **CaMK2a** | Calcium/calmodulin-dependent protein kinase type II alpha chain | VLAGQEYAAKIINTK | 60.9 | |
| **CaMK2b** | Calcium/calmodulin-dependent protein kinase type II beta chain | LCTGHEYAAKIINTK | 54.2 | |
| CaMK2d | Calcium/calmodulin-dependent protein kinase type II beta chain | IPTGQEYAAKIINTK | 25.1 | |
| CaMK2d | Calcium/calmodulin-dependent protein kinase type II beta chain | IPTGQEYAAKIINTKK | 25.4 | 22.7 |
| CaMK2g | Calcium/calmodulin-dependent protein kinase type II gamma chain | TSTQEYAAKIINTK | 50.6 | -6.5 |
| CaMK4 | Calcium/calmodulin-dependent protein kinase type IV | DLKPENLLYATPAPDAPLK | 29.6 | |
| CaMKK1 | Calcium/calmodulin-dependent protein kinase kinase 1 | LAYNESEDRHYAMKVLSK | 20.8 | |
| CaMKK2 | Calcium/calmodulin-dependent protein kinase kinase 2 | LAYNENDNTYYAMKVLSK | 15.3 | |
| CaMKK2 | Calcium/calmodulin-dependent protein kinase kinase 2 | DIKPSNLLVGEDGHIK | 8.5 | |
| CASK | Peripheral plasma membrane protein CASK | ETGQQFAVKIVDVAK | -21.3 | |
| CCRK | Cell cycle-related kinase | DLKPANLLISASGQLK | 20.9 | 16.2 |
| CDK2 | Cell division protein kinase 2 | DLKPQNLLINAEGSIK | | 9.5 |

| Kinase | Description | Sequence | Brain | Liver |
|---|---|---|---|---|
| **CDK11,CDK8** | Cell division protein kinase 8 | DLKPANILVMGEGPER | 66.8 | |
| CDK5 | Cell division protein kinase 5 | NRETHEIVALKR | 9.5 | 5.8 |
| CDK5 | Cell division protein kinase 5 | DLKPQNLLINR | 8.0 | |
| CDK5 | Cell division protein kinase 5 | NVLHRDLKPQNLLINR | 27.4 | |
| CDK7 | Cell division protein kinase 7 | DLKPNNLLLDENGVLK | | -21.8 |
| CHED | Cell division cycle 2-like protein kinase 5 | DIKCSNILLNNR | 1.1 | |
| CK1a | Casein kinase I isoform alpha | DIKPDNFLMGIGR | 36.8 | |
| CSK | Tyrosine-protein kinase CSK | VSDFGLTKEASSTQDTGK | -69.2 | |
| CSK | Tyrosine-protein kinase CSK | VSDFGLTKEASSTQDTGKLPVK | -13.0 | 2.8 |
| DCAMKL1 | Serine/threonine-protein kinase DCAMKL1 | DIKPENLLVYEHQDGSK | -7.6 | |
| DCAMKL1 | Serine/threonine-protein kinase DCAMKL1 | FSAVQVLEHPWVNDDGLPENEHQLSVAGKIK | -4.1 | |
| DCAMKL2 | Serine/threonine-protein kinase DCAMKL2 | DIKPENLLVCEYPDGTK | 14.3 | |
| eEF2K | Elongation factor 2 kinase | YIKYNSNSGFVR | -0.5 | -31.4 |
| EGFR | Epidermal growth factor receptor precursor | IPVAIKELR | -3.7 | -11.0 |
| EphA7 | Ephrin type-A receptor 7 precursor | VIEDDPEAVYTTTGGKIPVR | -15.4 | |
| EphB2 | Ephrin type-B receptor 2 precursor | FLEDDTSDPTYTSALGGKIPIR | -7.7 | |
| EphB3 | Ephrin type-B receptor 3 precursor | FLEDDPSDPTYTSSLGGKIPIR | 32.7 | |
| Erk1 | Mitogen-activated protein kinase 3 | DLKPSNLLINTTCDLK | -1.6 | 15.6 |
| Erk2 | Mitogen-activated protein kinase 14 | DLKPSNLLLNTTCDLK | | 17.3 |
| FAK | Focal adhesion kinase 1 | CIGEGQFGDVHQGVYLSPENPALAVAIKTCK | -25.0 | |
| FER | Proto-oncogene tyrosine-protein kinase FER | QEDGGVYSSSGLKQIPIK | 43.9 | |
| FER | Proto-oncogene tyrosine-protein kinase FER | DKTPVAIKTCKEDLPQELK | 45.6 | 10.0 |
| FER | Proto-oncogene tyrosine-protein kinase FER | TPVAIKTCKEDLPQELK | 46.4 | |
| FES | Proto-oncogene tyrosine-protein kinase Fes/Fps | LRADNTPVAVKSCR | -114.7 | 3.3 |
| FRAP (mTOR) | Serine/threonine-protein kinase mTOR | IQSIAPSLQVITSKQRPR | | -26.8 |
| FYN | Proto-oncogene tyrosine-protein kinase Fyn | VAIKTLKPGTMSPESFLEEAQIMKK | -12.4 | |
| FYN,SRC,YES | Proto-oncogene tyrosine-protein kinase Fyn | QGAKFPIKWTAPEAALYGR | 3.3 | |
| GCK | Mitogen-activated protein kinase kinase kinase kinase 2 | DTVTSELAAVKIVK | 18.8 | 4.6 |
| GCK | Mitogen-activated protein kinase kinase kinase kinase 2 | DIKGANLLLTLQGDVK | 33.1 | |
| GPRK4,GPRK6 | G protein-coupled receptor kinase 4 | DLKPENILLDDHGHIR | | 36.2 |
| GSK3A | Glycogen synthase kinase-3 alpha | DIKPQNLLVDPDTAVLK | 15.4 | 6.8 |
| GSK3B | Glycogen synthase kinase-3 beta | DIKPQNLLLDPDTAVLK | 38.8 | -2.1 |

| Kinase | Description | Sequence | Brain | Liver |
|---|---|---|---|---|
| HER4/ErbB4 | Receptor tyrosine-protein kinase erbB-4 | LLEGDEKEYNADGGKMPIK | 13.6 | |
| HER4/ErbB4 | Receptor tyrosine-protein kinase erbB-4 | GIWVPEGETVKIPVAIKILNET TGPK | 3.8 | |
| ILK | Integrin-linked protein kinase | GRWQGNDIVVKVLK | -22.1 | |
| ILK | Integrin-linked protein kinase | WQGNDIVVKVLK | -51.3 | 8.3 |
| ILK | Integrin-linked protein kinase | ISMADVKFSFQCPGR | -27.7 | |
| IRAK1 | Interleukin-1 receptor-associated kinase 1 [Mus musculus (Mouse)] | AIQFLHQDSPSLIHGDIKSSN VLLDER | | -47.8 |
| IRAK4 | Interleukin-1 receptor-associated kinase 4 | DIKSANILLDKDFTAK | 15.5 | -6.4 |
| JAK1 | Tyrosine-protein kinase JAK1 | IGDFGLTKAIETDKEYYTVK | | -7.6 |
| JAK1 | Tyrosine-protein kinase JAK1 | YDPEGDNTGEQVAVKSLKPE SGGNHIADLKK | | 13.6 |
| JAK1 domain1 | Tyrosine-protein kinase JAK1 | QLASALSYLEDKDLVHGNVC TKNLLLAR | | 25.4 |
| JNK1,JNK2,JNK3 | Mitogen-activated protein kinase 8 | DLKPSNIVVK | 3.0 | -24.8 |
| KHS1 | Mitogen-activated protein kinase kinase kinase kinase 5 | NVHTGELAAVKIIK | 1.3 | -13.0 |
| KHS1 | Mitogen-activated protein kinase kinase kinase kinase 5 | DIKGANILLTDHGDVK | 18.9 | |
| KHS2 | Mitogen-activated protein kinase kinase kinase kinase 3 | NVNTGELAAIKVIK | 10.6 | |
| LATS1 | Serine/threonine-protein kinase LATS1 | ALYATKTLR | | -9.5 |
| LKB1 | Serine/threonine-protein kinase LKB1 | DIKPGNLLLTTNGTLK | -8.5 | -3.4 |
| LOK | Serine/threonine-protein kinase 10 | DLKAGNVLMTLEGDIR | 18.6 | 15.2 |
| MAK | Serine/threonine-protein kinase MAK | SNESGELVAIKR | 4.8 | |
| MAP2K1 | Dual specificity mitogen-activated protein kinase kinase 1 | KLIHLEIKPAIR | | -7.6 |
| MAP2K1,MAP2K2 | Dual specificity mitogen-activated protein kinase kinase 1 | DVKPSNILVNSR | 14.0 | -26.2 |
| MAP2K2 | Dual specificity mitogen-activated protein kinase kinase 2 | KLIHLEIKPAVR | -5.0 | -14.0 |
| MAP2K3 | Dual specificity mitogen-activated protein kinase kinase 3 | DVKPSNVLINK | | -9.0 |
| MAP2K4 | Dual specificity mitogen-activated protein kinase kinase 4 | LCDFGISGQLVDSIAKTR | -1.3 | |
| MAP2K4 | Dual specificity mitogen-activated protein kinase kinase 4 | DIKPSNILLDR | 40.0 | 7.0 |
| **MAP2K6** | Dual specificity mitogen-activated protein kinase kinase 6 | DVKPSNVLINTLGQVK | 53.2 | -10.8 |
| MAP3K2 | Mitogen-activated protein kinase kinase kinase 2 [Mus musculus (Mouse)] | ELAVKQVQFNPESPETSKEV NALECEIQLLK | | 34.7 |
| MAP3K2,MAP3K3 | Mitogen-activated protein kinase kinase kinase 2 | DIKGANILR | -17.2 | -7.7 |
| MAP3K4 | Mitogen-activated protein kinase kinase kinase 4 | DIKGANIFLTSSGLIK | | -34.7 |
| MAP3K5 | Mitogen-activated protein kinase kinase kinase 5 | DIKGDNVLINTYSGVLK | 20.3 | 9.3 |
| MAP3K5,MAP3K6 | Mitogen-activated protein kinase kinase kinase 5 | IAIKEIPER | 5.2 | |
| MAP3K5,MAP3K6 | Mitogen-activated protein kinase kinase kinase 5 | IAIKEIPERDSR | | 15.3 |
| MARK1 | MAP/microtubule affinity-regulating kinase 3 | EVAVKIIDKTQLNPTSLQK | 8.5 | |
| MARK2 | Serine/threonine-protein kinase MARK2 | EVAVKIIDKTQLNSSSLQK | -20.9 | |

| Kinase | Description | Sequence | Brain | Liver |
|---|---|---|---|---|
| MARK2 | Serine/threonine-protein kinase MARK2 | HILTGKEVAVKIIDKTQLNSSSLQK | -6.9 | |
| MARK3 | MAP/microtubule affinity-regulating kinase 3 | EVAIKIIDKTQLNPTSLQK | -2.2 | |
| MARK3,MARK4 | MAP/microtubule affinity-regulating kinase 3 | EVAIKIIDK | -9.4 | |
| MARK4 | MAP/microtubule affinity-regulating kinase 4 | EVAIKIIDKTQLNPSSLQK | 23.2 | |
| MER,TYRO3 | Proto-oncogene tyrosine-protein kinase MER precursor | KIYSGDYYR | 22.6 | -21.7 |
| MET | Hepatocyte growth factor receptor precursor | DMYDKEYYSVHNK | | -16.8 |
| MLK1 | Mitogen-activated protein kinase kinase kinase 9 | DLKSSNILILQK | 6.4 | |
| MLK2 | Mitogen-activated protein kinase kinase kinase 10 | DLKSINILILEAIENHNLADTVLK | -14.9 | |
| MST1 | STE20-like kinase MST1 | ETGQIVAIKQVPVESDLQEIIK | -28.6 | |
| MST1,MST2 | STE20-like kinase MST1 | DIKAGNILLNTEGHAK | -18.6 | -2.3 |
| MST2 | Serine/threonine-protein kinase 3 | ESGQVVAIKQVPVESDLQEIIK | -36.1 | 5.3 |
| MST3 | Serine/threonine-protein kinase 24 | DIKAANVLLSEHGEVK | 1.0 | 3.2 |
| MST3,MST4,YSK1 | Serine/threonine-protein kinase 24 | LADFGVAGQLTDTQIKR | 9.1 | |
| MST4,YSK1 | Serine/threonine-protein kinase 25 | DIKAANVLLSEQGDVK | -55.8 | -31.5 |
| NDR1 | Serine/threonine-protein kinase 38 | DTGHVYAMKILR | -5.3 | 29.6 |
| NDR1,NDR2 | Serine/threonine-protein kinase 38 | LSDFGLCTGLKK | -7.5 | |
| NDR2 | Serine/threonine-protein kinase 38-like | DTGHIYAMKILR | 36.7 | 10.7 |
| NEK1 | Serine/threonine-protein kinase Nek1 | DIKSQNIFLTK | 13.9 | |
| NEK4 | Serine/threonine-protein kinase Nek4 | DLKTQNVFLTR | | -21.9 |
| NEK7 | Serine/threonine-protein kinase Nek7 | DIKPANVFITATGVVK | -24.9 | 1.3 |
| NIM1 | Hypothetical eukaryotic protein kinase containing protein | VAIKILDK | -23.9 | |
| NuaK1 | NUAK family, SNF1-like kinase 1 | VVAIKSIR | 2.9 | |
| OSR1 | Serine/threonine-protein kinase OSR1 | DVKAGNILLGEDGSVQIADFGVSAFLATGGDITR | -20.5 | -70.0 |
| p38a | Mitogen-activated protein kinase 14 | QELNKTIWEVPER | 5.5 | 31.9 |
| p38d,p38g | Mitogen-activated protein kinase 12 | DLKPGNLAVNEDCELK | | -8.1 |
| p70S6K | Ribosomal protein S6 kinase I | DLKPENIMLNHQGHVK | -6.1 | 5.9 |
| p70S6K,p70S6Kb | Ribosomal protein S6 kinase I | GGYGKVFQVR | -12.0 | |
| PCTAIRE1,PCTAIRE2,PCTAIRE3 | Serine/threonine-protein kinase PCTAIRE-1 | DLKPQNLLINER | -3.3 | -25.4 |
| PCTAIRE2,PCTAIRE3 | Serine/threonine-protein kinase PCTAIRE-2 | SKLTENLVALKEIR | -1.6 | |
| PFTAIRE1 | Serine/threonine-protein kinase PFTAIRE-1 | LVALKVIR | 18.0 | |
| PFTAIRE1 | Serine/threonine-protein kinase PFTAIRE-1 | DLKPQNLLISDTGELK | 19.7 | |

| Kinase | Description | Sequence | Brain | Liver |
|---|---|---|---|---|
| PHKg1 | Phosphorylase b kinase gamma catalytic chain, skeletal muscle isoform | CIHKPTCQEYAVKIIDITGGGS FSSEEVQELR | -8.4 | |
| PI4KA | Phosphatidylinositol 4-kinase, catalytic, alpha polypeptide | SGTPMQSAAKAPYLAK | 39.9 | 21.7 |
| PI4KB | Phosphatidylinositol 4-kinase beta | VPHTQAVVLNSKDK | -2.1 | -18.6 |
| PI4KB | Phosphatidylinositol 4-kinase beta | VPHTQAVVLNSKDKAPYLIYV EVLECENFDTTSVPARIPENR | 23.1 | |
| PIP4K2A | Phosphatidylinositol-4-phosphate 5-kinase type II alpha | AKELPTLKDNDFINEGQK | -11.3 | |
| PIP4K2A | Phosphatidylinositol-4-phosphate 5-kinase type II alpha | ELPTLKDNDFINEGQK | -27.7 | |
| PIP4K2B | Phosphatidylinositol-4-phosphate 5-kinase type II beta | AKDLPTFKDNDFLNEGQK | -20.0 | |
| PIP4K2B | Phosphatidylinositol-4-phosphate 5-kinase type II beta | DLPTFKDNDFLNEGQK | -18.4 | |
| PIP4K2C | Phosphatidyl inositol phosphate kinase type II gamma | TLVIKEVSSEDIADMHSNLSN YHQYIVK | 28.8 | |
| PIP4K2C | Phosphatidyl inositol phosphate kinase type II gamma | VKELPTLKDMDFLNK | 37.2 | |
| PIP5K3 | FYVE finger-containing phosphoinositide kinase | GGKSGAAFYATEDDRFILK | 16.3 | 0.6 |
| PITSLRE | PITSLRE serine/threonine-protein kinase CDC2L1 | DLKTSNLLLSHAGILK | 31.2 | |
| PKACa | cAMP-dependent protein kinase catalytic subunit alpha | DLKPENLLIDQQGYIQVTDFG FAK | 13.8 | |
| PKCe | Protein kinase C epsilon type | DLKLDNILLDAEGHCK | -24.1 | |
| PKR | Interferon-induced, double-stranded RNA-activated protein kinase | DLKPGNIFLVDER | 31.5 | -7.1 |
| PLK1 | Serine/threonine-protein kinase PLK1 | CFEISDADTKEVFAGKIVPK | | 7.9 |
| QSK | Serine/threonine-protein kinase QSK | VAIKIIDKSQLDEENLKK | -7.4 | |
| QSK | Serine/threonine-protein kinase QSK | DLKAENLLLDANLNIK | -12.8 | |
| RIPK2 | Receptor-interacting serine/threonine-protein kinase 2 | ILHEIALGVNYLHNMNPPLLH HDLKTQNILLDNEFHVK | | -4.3 |
| RIPK3 | Receptor-interacting serine/threonine-protein kinase 3 | DLKPSNILLDPELHAK | | -1.7 |
| ROCK1 | Rho-associated protein kinase 1 | KLQLELNQER | 0.0 | -9.6 |
| ROCK1,ROCK2 | Rho-associated protein kinase 2 | DVKPDNMLLDK | 41.6 | |
| RSK1 domain1 | Ribosomal protein S6 kinase alpha 1 | DLKPENILLDEEGHIKLTDFGL SKEAIDHEK | | -7.4 |
| RSK1 domain2 | Ribosomal protein S6 kinase alpha 2 | DLKPSNILYVDESGNPECLR | -29.0 | 16.1 |
| RSK1,RSK2,RSK 3 domain1 | Ribosomal protein S6 kinase alpha 1 | DLKPENILLDEEGHIK | -3.4 | |
| RSK1,RSK3 domain2 | Ribosomal protein S6 kinase alpha 1 | SKRDPSEEIEILLR | -68.1 | |
| RSK2 domain1 | Ribosomal protein S6 kinase alpha 3 | DLKPENILLDEEGHIKLTDFGL SKESIDHEK | | -56.2 |
| **Kinase** | **Description** | **Sequence** | **Brain** | **Liver** |

| Kinase | Description | Sequence | Brain | Liver |
|---|---|---|---|---|
| RSK3 domain1 | Ribosomal protein S6 kinase alpha 1 | DLKPENILLDEEGHIKITDFGLSK | 3.6 | 19.2 |
| RSK3 domain2 | Ribosomal protein S6 kinase alpha 1 | DLKPSNILYMDESGNPESIR | -20.5 | |
| RSKL1 domain1 | Ribosomal protein S6 kinase delta-1 | VLGVIDKVLLVMDTR | -10.2 | |
| RSKL1 domain1 | Ribosomal protein S6 kinase delta-1 | VLGVIDKVLLVMDTRTEQTFILK | 5.7 | |
| SGK2 | Serine/threonine-protein kinase Sgk2 | KSDGAFYAVKVLQK | | 14.9 |
| SGK3 | Serine/threonine-protein kinase Sgk3 | FYAVKVLQK | -29.0 | |
| SLK | STE20-like serine/threonine-protein kinase | ETNVLAAAKVIDTK | 9.9 | |
| SLK | STE20-like serine/threonine-protein kinase | DLKAGNILFTLDGDIK | -17.7 | -24.3 |
| SLK | STE20-like serine/threonine-protein kinase | DLKAGNILFTLDGDIKLADFGVSAK | 9.8 | |
| SLK | STE20-like serine/threonine-protein kinase | IIHRDLKAGNILFTLDGDIK | 18.9 | |
| SMG1 | Serine/threonine-protein kinase SMG1 | DTVTIHSVGGTITILPTKTKPK | 18.7 | -2.6 |
| SMG1 | Serine/threonine-protein kinase SMG1 | SYPYLFKGLEDLHLDER | | -28.8 |
| SNRK | SNF-related serine/threonine-protein kinase | DLKPENVVFFEK | -3.8 | |
| SRPK1,SRPK2 | Serine/threonine-protein kinase SRPK1 | FVAMKVVK | -8.0 | |
| STLK5 | Protein kinase LYK5 splice variant 1 homolog | AVHDFPKYSIKVLPWLSPEVLQQNLQGYDAK | 21.4 | |
| STLK5 | Protein kinase LYK5 splice variant 1 homolog | YSIKVLPWLSPEVLQQNLQGYDAK | 7.8 | 37.8 |
| STLK6 | Serine/threonine-protein kinase ALS2CR2 | SFKASHILISGDGLVTLSGLSHLHSLLK | -3.0 | |
| TAK1 | Mitogen-activated protein kinase kinase kinase 7 | DLKPPNLLLVAGGTVLK | -3.3 | |
| TAO1,TAO3 | Serine/threonine-protein kinase TAO3 | DIKAGNILLTEPGQVK | 7.4 | -38.1 |
| TAO2 | Serine/threonine-protein kinase TAO2 | DVKAGNILLSEPGLVK | 50.9 | -0.7 |
| TLK1 | Serine/threonine-protein kinase tousled-like 1 | YLNEIKPPIIHYDLKPGNILLVDGTACGEIK | 46.6 | 16.5 |
| TLK1,TLK2 | Serine/threonine-protein kinase tousled-like 1 | YLNEIKPPIIHYDLKPGNILLVNGTACGEIK | | 11.8 |
| ULK3 | hypothetical MIT/Serine/threonine protein kinase containing protein | EVVAIKCVAK | 37.5 | |
| ULK3 | hypothetical MIT/Serine/threonine protein kinase containing protein | NISHLDLKPQNILLSSLEKPHLK | | 34.8 |
| VACAMKL | CaM kinase-like vesicle-associated protein | NYNQPSEVTDRYDLGQVIKTEEFCEIFR | -40.4 | |
| Wnk1,Wnk2,Wnk4 | Serine/threonine-protein kinase WNK2 | IGDLGLATLKR | 12.5 | |

| | | | | |
|---|---|---|---|---|
| YANK3 | Serine/threonine kinase 32C | DVKPDNILLDEQGHAHLTDF NIATIIK | -17.5 | |
| ZC1/HGK,ZC2/TNIK,ZC3/MINK | Mitogen-activated protein kinase kinase kinase kinase 4 | DIKGQNVLLTENAEVK | -17.0 | -11.9 |
| ZC2/TNIK | Traf2 and NCK-interacting protein kinase | TGQLAAIKVMDVTGDEEEEIK QEINMLK | -63.8 | |
| ZC2/TNIK | Traf2 and NCK-interacting protein kinase | TGQLAAIKVMDVTGDEEEEIK QEINMLKK | 4.7 | |
| ZC3/MINK | Misshapen-like kinase 1 | TGQLAAIKVMDVTEDEEEEIK QEINMLKK | -10.8 | |
